# Real time, field-deployable whole genome sequencing of malaria parasites using nanopore technology

**DOI:** 10.1101/2020.12.17.423341

**Authors:** Zahra Razook, Somya Mehra, Brittany Gilchrist, Digjaya Utama, Dulcie Lautu-Gumal, Abebe Fola, Didier Menard, James Kazura, Moses Laman, Ivo Mueller, Leanne J. Robinson, Melanie Bahlo, Alyssa E. Barry

## Abstract

Malaria parasite genomes have been generated predominantly using short read sequencing technology which can be slow, requires advanced laboratory training and does not adequately interrogate complex genomic regions that harbour important malaria virulence determinants. The portable Oxford Nanopore Technologies MinION platform generates long reads in real time and may overcome these limitations. We present compelling evidence that Nanopore sequencing delivers valuable additional information for malaria parasites with comparable data fidelity for single nucleotide variant (SNV) calls, compared to standard Illumina whole genome sequencing. We demonstrate this through sequencing of pure *Plasmodium falciparum* DNA, mock infections and natural isolates. Nanopore has low error rates for haploid SNV genotyping and identifies structural variants (SVs) not detected with short reads. Nanopore genomes are directly comparable to publically available genomes and produce high quality end to end chromosome assemblies. Nanopore sequencing will expedite genomic surveillance of malaria and provide new insights into parasite genome biology.

## INTRODUCTION

Whole genome sequencing (WGS) provides complete information about pathogens that along with epidemiological metadata can enhance control efforts^1^ to track the spread of infectious diseases in real time^2^, the emergence of drug resistance^3^, responses to control interventions^4^ and to inform vaccine design^5^. Human malaria, a disease that has plagued humans for thousands of years, is caused by infection with *Plasmodium* species, the most virulent of which is *Plasmodium falciparum*. Malaria remains one of the world’s most widespread and deadly infectious diseases killing more than 400 people annually, and causing over 200 million clinical episodes of the disease^6^. A major roadblock to WGS of clinical (or “field”) isolates has recently been overcome through methods to enrich trace amounts of parasite DNA from finger prick blood samples contaminated with large amounts of human DNA^7, 8^. However, generation of WGS has traditionally been restricted to well-equipped laboratories that can maintain large, expensive sequencing platforms as well as advanced analytical pipelines and human resources required for data processing. As Illumina short read WGS (srWGS) requires extensive human expertise in both library preparation and instrument support, researchers in malaria-endemic countries have generally needed to send samples to large genome centres for sequencing resulting in significant time delays and loss of data custodianship. However for genomic surveillance to inform malaria control and elimination in a timely manner, large numbers of genomes with spatially dense sampling and the rapid generation of high quality data at low cost will be needed^9^ and this is only feasible through sequencing in the field.

The Oxford Nanopore Technologies (ONT) MinION is a portable, pocket-sized device that operates through a USB port in a personal computer. Nanopore sequencing involves the movement of DNA through a synthetic porous membrane (flow cell) resulting in unique ionic currents for each six basepair window that are then translated into nucleotides in real time. The MinION device is an attractive option for malaria genomic surveillance because it can be deployed to field or clinic settings with minimal capital cost and training, and thus provides potential for rapid genomic profiling. In the clinical setting, MinION would provide a platform to rapidly screen for multigene haplotypes associated with drug resistance^10, 11^, and may provide a pathway for personalised treatment in some well-resourced settings^12^. Long read WGS (lrWGS) using MinION could be used to rapidly track the origins of outbreaks as has been the case for other human pathogens^2, 13^. The low cost, relative portability and ease of use compared to other platforms suggests this platform has utility for both public health surveillance and capacity strengthening^14^. Data can be collected for future studies and can continue to be analysed for a variety of additional features as knowledge expands.

Unique features of the *Plasmodium falciparum* genome contribute to the challenge of WGS. The genome is AT-rich (81%), with extended tracts of repetitive, low-complexity DNA^15^, particularly in subtelomeric regions. Hypervariable, multigene families, including the *var, rifin* and *stevor* families, have been difficult to characterise with srWGS because divergent short reads cannot be reliably aligned to a reference genome, although specialised pipelines have been developed^16^. In addition, malaria infections are often multiclonal, especially in high transmission regions^17^ and therefore an added challenge for malaria population genomics is reconstructing these clonal haplotypes. LrWGS has the potential to overcome these limitations by spanning complex regions, to enable more accurate assembly and to differentiate clonal haplotypes within mutliple infections.

We aimed to develop and optimise Nanopore sequencing protocols for *P. falciparum* and to benchmark the resulting lrWGS against that obtained using the current gold standard Illumina srWGS. Initially, we tested the platform on pure *P. falciparum* DNA extracted from cultured strains, and compared the data to publically available reference genomes. We then optimised the sequencing protocol for natural human infections using mock infections comprising DNA from a reference *P. falciparum* strain spiked with human DNA. Finally, we performed shallow sequencing of a number of field isolates from Papua New Guinea (PNG) and Cambodia, allowing key parameters for field sample sequencing to be determined. The resulting data was used to estimate baseline error rates, to quantify the accuracy of single nucleotide variant (SNV) genotypes, to obtain drug resistance profiles, and to benchmark against publically available Illumina srWGS data from the same countries. Deep sequencing of cultured strains allowed *de novo* assemblies. These high quality reference genomes enable characterisation of full length *var* genes and mapping of large structural variants (SVs), which were not achievable with srWGS. Optimised sequencing protocols and pipelines are provided so that this approach can be implemented in other laboratories to permit true ‘sequencing in the field’.

## RESULTS

### Nanopore sequencing of *P. falciparum*

In order to test the capability of MinION for sequencing *P. falciparum*, test runs were first performed using pure *P. falciparum* DNA from the XHA^18^ and BB12 (a descendant of the Brazilian IT isolate^19, 20^) cultured lines with each run on an independent flow cell for the full 48 hours recommended by ONT. The total output of each run was 3.42 and 2.07 Gb, with N50 read lengths of 14.63 and 15.26 kB (mean read lengths of 8.80 and 6.95 kb) and maximum read lengths of 261.32 and 157.32 kb respectively (Table S2A), with quality scores of more than Q5 for over 90% of reads. The two samples produced 120X and 76X mean depth of coverage (reads), with 95% of the genome covered by more than 30 reads and relatively uniform genome-wide coverage (Figure 1A).

**Figure 1.**
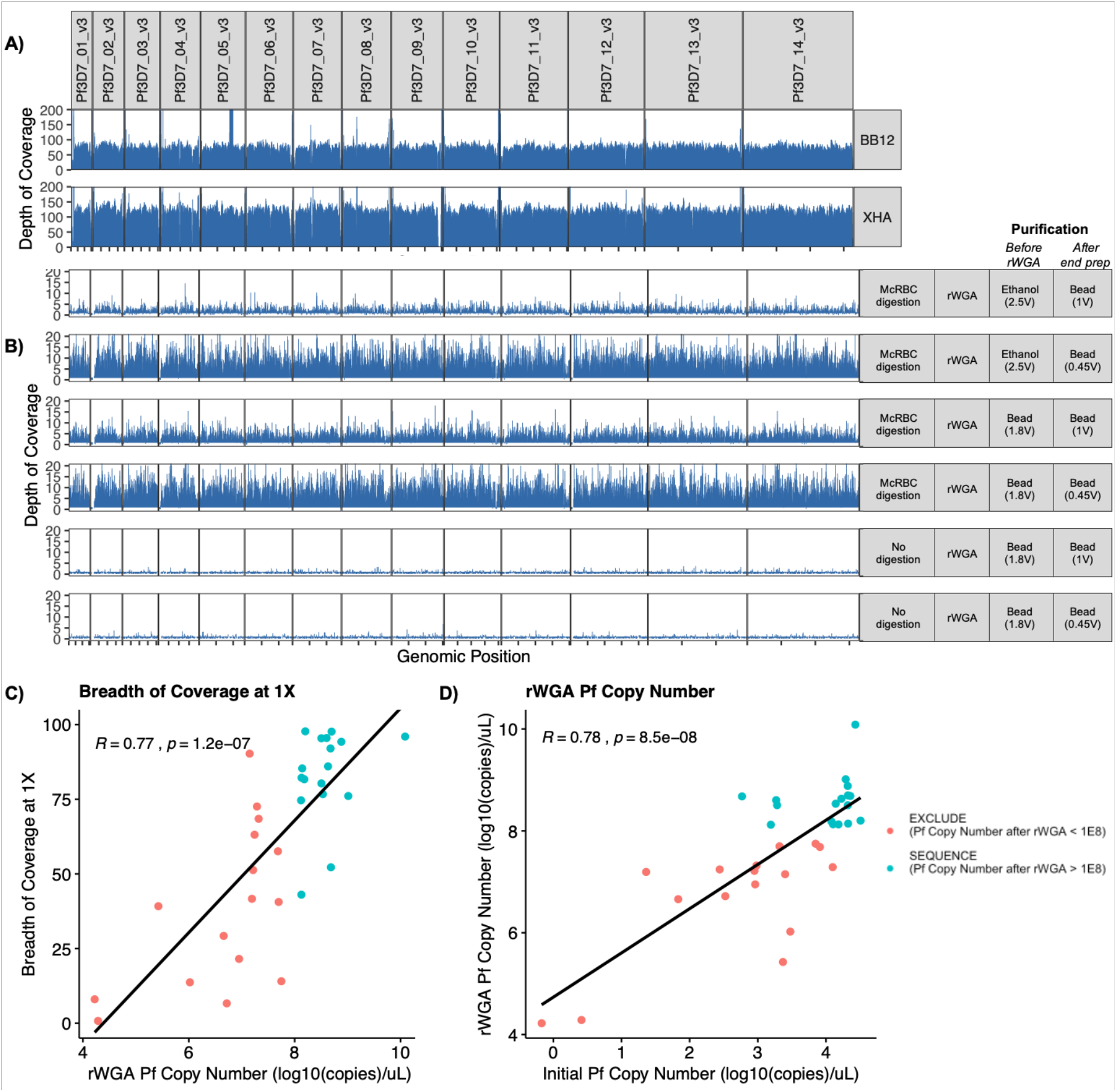
Output of MinION sequencing runs. **(A)** Schematic of genome coverage assessed in 5kb bins of the parasite genome for pure P. falciparum clones (cultured lines). Coverage is a term that can be used interchangeably to describe both depth and breadth of genome coverage. Depth refers to the number of times a base in the reference genome is covered by a read and correlates to greater confidence in variant calling. Breadth refers to the percentage of the target genome that is covered by reads in a sequencing experiment. High uniformity, breadth and depth of coverage were attained for both BB12 and XHA. **(B)** Optimisation of parasite DNA amplification and library preparation using mock infections. Samples comprising 1% P. falciparum DNA (3D7) were spiked with human DNA and subject to different enrichment (digestion with McrBC and/or rWGA). The depth of sequencing coverage across the parasite genome is shown for each condition. Sequencing was run in multiplex for the six conditions. **(C)** Log-transformed post-enrichment P. falciparum 18s rRNA gene copy numbers (as a measure of Pf DNA concentration within a sample) are most strongly correlated with the breadth of coverage at 1X (the proportion of the genome covered by at least one read). A Pf copy number threshold of with 1.0×10^8^ is the most appropriate diagnostic for a minimum breadth of coverage of 75% at 1X based on a receiver-operator characteristic analysis (data not shown). **(D)** rWGA Pf copy numbers after enrichment as a function of the pre-enrichment (initial) Pf copy number. Datapoints are coloured on the basis of whether they met the criteria for exclusion (red) or sequencing (green). Based on these findings we recommend selecting field isolates with a minimum of 1000 parasite copies/μL for enrichment, and enriched samples with at least 1.0×10^8^ parasite copies/μL for sequencing.

*P. falciparum* field isolates obtained by collecting the blood of human volunteers contain large amounts of contaminating human DNA after DNA extraction^21^. Mock infections of 1% parasitemia (20,000 parasites/μL^8^) were subject to different enrichment conditions. The resulting six samples were run in multiplex on a single 48hr run. The enrichment procedure (*McrBC* digest plus random whole genome amplification (rWGA)) significantly increased the breadth and depth of parasite genome coverage (Figure 1B, Table S2B) and the proportion of parasite DNA (≥83%) compared to rWGA alone (≤5%). After end repair, the 0.45V bead cleanup was associated with higher genome-wide read coverage than 1V bead clean-up. However, both 2.5V ethanol precipitation and 1.8V SPRISelect bead purification prior to library preparation performed similarly in terms of the read quality and output (Table S2B).

Using the optimised Nanopore sequencing protocol (1.8V bead purification after enrichment and 0.45V beads purification after end prep), we then sequenced 33 *P. falciparum* field isolates, multiplexing three to four samples per 48 hour run (Table S2C). Field samples ranged in starting parasite densities from 0.674–32,200 18s rRNA copies/uL (Q1=916 copies/uL, Median=3,000 copies/uL, Q3=16,840 copies/uL). Methylation-dependent *McrBC* digestion and whole genome amplification resulted in between 114–810,000 fold enrichment (Q1=7,940, Median=21,200, Q3=52,900). The proportion of reads mapping to the parasite genome varied between 0% and 85% (Q1=6%, Median=18%, Q3=36%). Median read lengths ranged from 990–2,790 bp.

Key parameters associated with sequencing quality for individual field isolates are shown in Figure 1C-D. After enrichment, parasite DNA concentration (as measured by 18s rRNA copy number per μL) exhibited the strongest correlation with the breadth of genome wide coverage at 1X (R=0.77, p=1.2E-7) (Figure 1C). A copy number threshold of 1.0×10^8^ was found to be the most appropriate diagnostic for ≥75% breadth of coverage at 1X in a receiver-operating characteristic curve analysis (data not shown). Based on these results, enriched samples with a minimum of 1.0×10^8^ copies/μL are recommended for successful sequencing. A comparison of copy numbers before and after enrichment (Figure 1D) suggests that DNA samples with a minimum of 1000 copies/μL should be selected for enrichment.

Stratification by flow cell showed an association between the sequencing output and the amount of DNA loaded into a flowcell (Figure S2). A general, decreasing trend between the total number of bases sequenced and the amount of DNA loaded was evident above 600ng. Similarly, the proportion of reads with quality scores above 10 decreased as the amount of DNA loaded into a flowcell increased. Hence, clogged pores, were more likely when higher amounts of DNA (>600 ng) were loaded into the flowcell.

### Baseline sequencing error rates in Nanopore and Illumina sequencing

To quantify baseline error rates for Nanopore and Illumina, we compared 3D7 WGS to the publicly available *P. falciparum* 3D7 reference genome (version 3). While the core *P. falciparum* genome is stable in long-term *in vitro* culture, mitotic recombination can drive variation in antigen genes and subtelomeric regions^22^. We therefore restricted the reference genome-based analyses to SNVs situated in the core genome, with reference alleles to be treated as truth calls and alternate alleles as error. We focused on two key use cases for SNV calling: *de novo* SNV discovery (e.g. to interrogate novel variants associated with drug resistance) and SNV genotyping at a set of known loci (e.g. to conduct population genomic analyses or drug resistance profiling).

First, we considered false discovery rates (FDRs) for the detection of haploid *de novo* SNVs using each platform focusing on coding SNVs in 1356 “housekeeping” genes essential for *in vitro* asexual blood stage development^23^, with a cumulative transcript length of 2,792,980 bp. Illumina detected no coding SNVs in these genes relative to the reference genome while Nanopore sequencing data gave rise to 15 coding SNVs after relatively lenient filtration (i.e. minimum depth of coverage 5X, with at least 75% of reads supporting the called allele), with seven SNVs retained when the minimum depth of coverage was increased to 10X. These results suggest a higher FDR for *de novo* SNV discovery using Nanopore data relative to Illumina data, though it remains almost negligibly low. Albeit low coverage will also lead to lower power to detect novel variants. FDRs are contingent on coverage and variant filtration parameters - unlike the Illumina-specific GATK pipeline which considers well-validated metrics to filter out poor alignments, best practices workflows are yet to be developed for Nanopore SNV genotyping.

Next, baseline error rates for Nanopore and Illumina genotypes at a set of 742,365 validated SNV loci were analysed. Of 684,737 SNVs successfully genotyped by both platforms, Illumina incorrectly called 13 alternate alleles, while Nanopore incorrectly called 40 alternate alleles after minimal filtration (i.e. minimum depth of coverage 2X, with at least 75% of reads supporting the called allele). Baseline error rates for genotyping SNVs at validated loci therefore seem comparable for Nanopore and Illumina and appear to be very low across both platforms (<0.006%). However, since we have mapped reads against an isogenic reference genome, read alignments are likely to be correct and high genotyping accuracy may be driven by reference bias. It is therefore important to perform additional benchmarking of Nanopore and Illumina genotypes for isolates for which there is no isogenic reference genome.

### Accuracy of Nanopore genotypes

WGS analyses of *P. falciparum* commonly utilize high quality SNV calls to conduct population genomic analyses^24^. To validate the genotyping accuracy of Nanopore and analysis pipeline relative to Illumina, we performed a comparison of SNV genotypes derived from strains BB12 (Brazil) and XHA (Papua New Guinea, PNG) which have very high coverage (>70X), and two Cambodian field isolates with low coverage (<5X) for which we also collected Illumina data. Data was aligned to the 3D7 reference genome (version 3) and variants called as per 3D7 analysis. We then considered allele calls at 742,635 high-quality variants obtained through an in-house re-analysis of the srWGS MalariaGEN Pf3k plus PNG dataset^24, 25^ comprising 2,661 field isolates sampled from 15 malaria endemic countries. Only haploid genotypes determined by calling a single dominant allele at each locus were considered.

Nanopore SNV calls exhibited strong bias towards reference allele calls at very low depths of coverage. Moreover, at low depth, alternate (non-reference) alleles were generally called as reference alleles by the Nanopore analysis pipeline when the proportion of reads supporting the called genotype was relatively low, suggestive of conservative allele calls. Reference alignment bias, which arises during read mapping, has previously been reported for Nanopore (MinION) SNV genotyping^26^, with implications for haplotype reconstruction due to the overrepresentation of reference haplotypes^27^.

Since Nanopore sequencing has been shown to exhibit significant systematic error in homopolymeric regions^28^ we also stratified allele calls based on genomic content, by differentiating between calls within homopolymeric tracts of length greater than 6 bp and those in non-homopolymeric regions. Alignments around discordant calls in homopolymeric tracts were poor, with lower coverage than flanking genomic regions and a smaller proportion of reads supporting the called allele (Figure 2). In non-homopolymeric regions, discordant calls exhibited similar signatures, with a lower proportion of reads mapping to the called allele, but less substantial reductions in coverage. Discordant calls between Nanopore and Illumina could thus be diagnosed on the basis of low coverage and a low proportion of reads mapping to the called allele (that is, false heterozygosity) (Figure 2). Poor alignments of Nanopore sequence data around homopolymer tracts^28^ and false heterozygosity due to Nanopore sequencing error, particularly at low depths of coverage^29^, have been previously documented.

**Figure 2.**
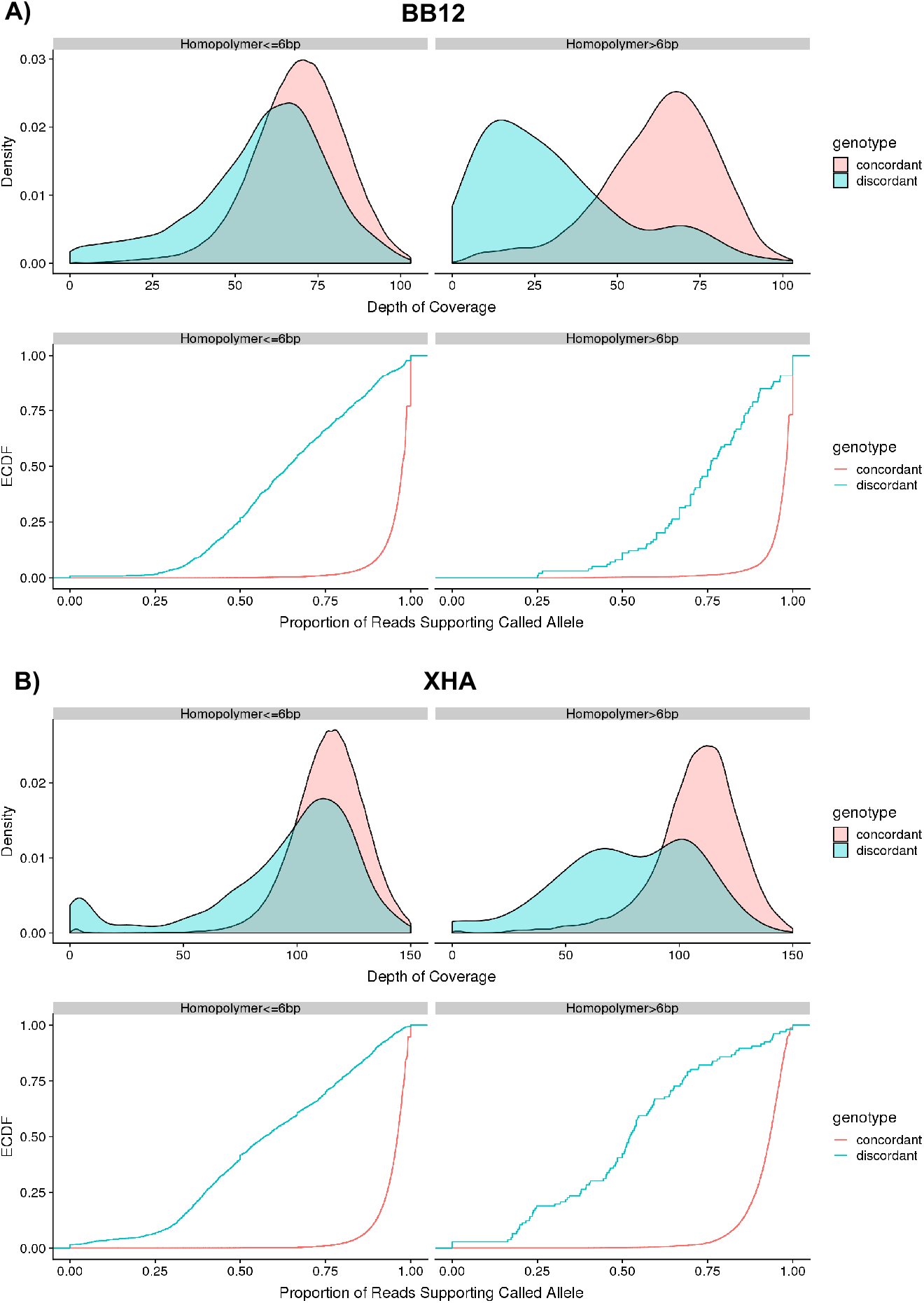
Variant quality metrics for discordant and concordant MinION genotype calls in both homopolymeric (>6bp) and non-homopolymeric regions for (A) BB12 and (B) XHA. Density plots show distributions for the depth of coverage by locus. Empirical cumulative distribution functions (ECDFs) show distributions for the proportion of reads supporting the called allele. In both homopolymeric and non-homopolymeric regions, discordant calls frequently had lower depths of coverage and a smaller proportion of reads supporting the called allele than concordant calls.

As a strategy to remove artefacts, loci were filtered by the depth of coverage and the proportion of reads supporting the allele call; previous studies have adopted these metrics^10^. Excluding loci with read support across multiple alleles does not account for heterozygosity or multiclonality. However, this approach was deemed appropriate since the cultured strains were presumed to be monoclonal, and shallow sequencing of field isolates generated inadequate coverage to detect minor clones. For field isolates (Table S2C) we only retained those genotypes with a minimum depth of coverage 2X where at least 75% of reads supported the called genotype (i.e. if 2X coverage, both reads must support the call).

Overall discordance rates between haploid Nanopore and lllumina genotypes were consistently low, even for Cambodian field isolates with poor mean coverage (less than 4X) however these isolates also had a large proportion of missing loci after filtering (Table 1). Nanopore reference calls had accuracy above 99.9% for all four isolates (Table 1). However, alternate allele calls were more error prone even after filtration, with only 88% supported by Illumina for Cam_01 (3.2X coverage) and Cam_02 (1.4X coverage). For cultured lines BB12 (75X coverage) and XHA (106X coverage) however, over 95% of Nanopore alternate allele calls were supported by Illumina calls. These results indicate that Nanopore alternate allele calls are significantly less reliable at low depths of coverage. As expected, discordance rates in homopolymeric tracts greater than 6 bp were found to be substantially higher than overall discordance rates (Table S3), highlighting the need to exercise caution in analysis of calls from homopolymeric tracts.

**Table 1:**
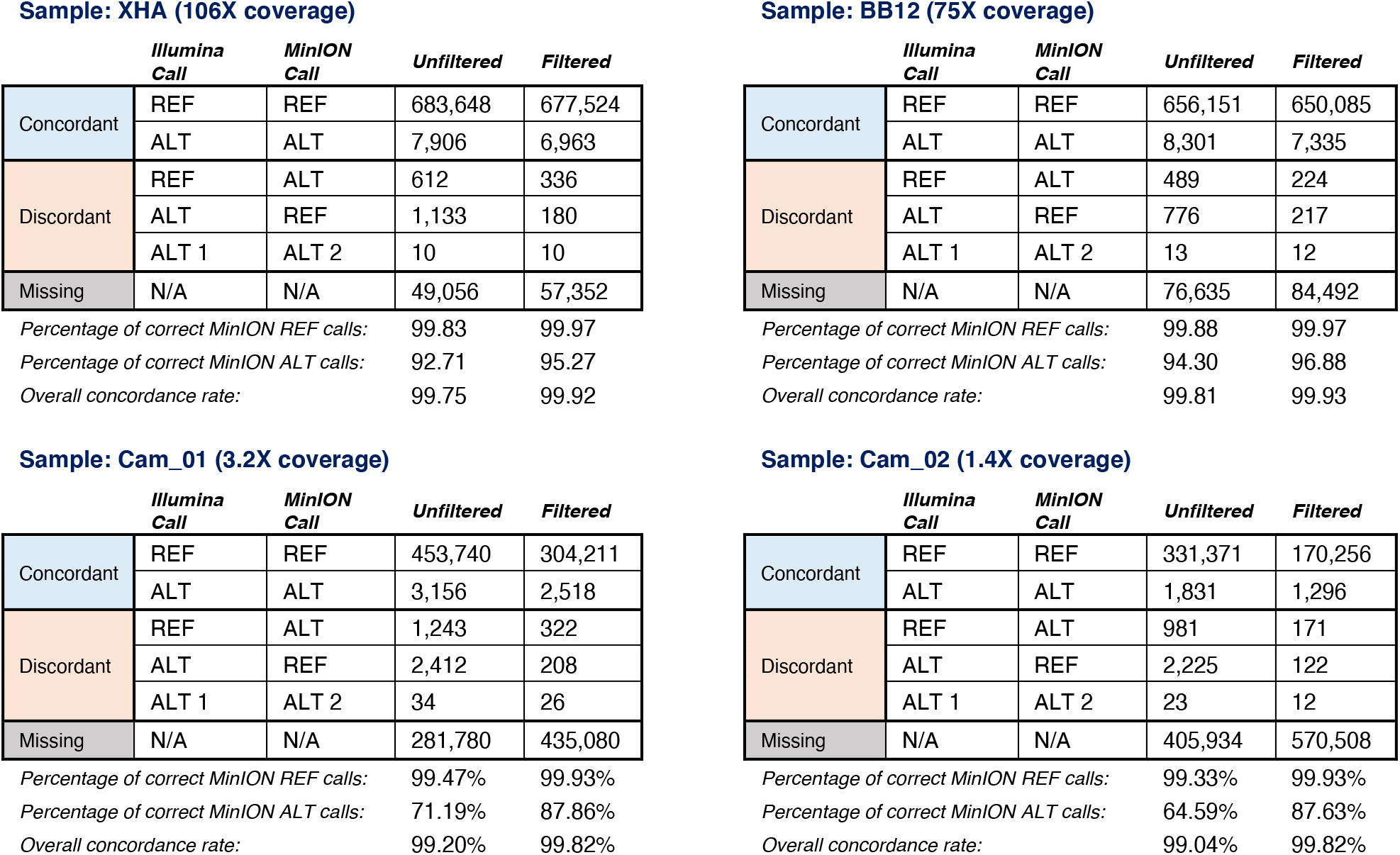
Comparison of Haploid MinION and Illumina Genotypes at 742,365 High-Quality Loci

Random errors in Nanopore sequencing reads were mitigated by increasing coverage, as evidenced by the lower discordance rate for isolates with high coverage. However, the persistence of genotyping error at high depths of coverage points to systematic differences between the two platforms. We therefore attempted to identify characteristic features of concordant and discordant calls that passed quality filtration.

The relative frequencies of concordant and discordant genotype calls after filtration were stratified by the variation type at discordant sites found within Nanopore and Illumina data. The most common discordant genotypes across the trialled isolates were the Illumina (treated to be truth calls) → Nanopore (treated as erroneous calls) transitions G → A and C → T, followed by the transversions T → A and A → T (Figure S3). In general, transition errors between similarly-structured bases (i.e. between two-ring purines or one-ring pyrmidines) were more pronounced than transversion errors for Nanopore sequencing, which might reflect errors in the base calling algorithm. Given that Nanopore calls are generated based on minute changes in current as the DNA passes through the pore, it is not surprising that biochemically similar bases are harder to distinguish and result in erroneous calls.

### Drug resistance profiling using Nanopore sequencing

As a measure of the capability of Nanopore sequencing to produce functionally relevant information from skim sequencing of field isolates, we determined SNV genotypes for each field isolate across a range of known drug resistance marker loci including *crt* (chloroquine, amodiaquine, piperaquine), *dhfr-ts* (pyrimethamine), *dhps* (sulfadoxine), *mdr1* (lumefantrine, mefloquine) and *kelch13* (artemisinin). Only genotype calls with depth of coverage at least 2X and at least 75% of reads supporting the called allele were retained. Functional annotations were applied to filtered variants, allowing the extraction of both nucleotide and amino acid haplotypes.

Haplotypes generated from Nanopore and Illumina sequencing data for the cultured lines BB12 and XHA, and field isolate Cam_01 were compared first. Filtered drug resistance gene haplotypes were generally concordant between the two sequencing platforms, with the exception of CRT codons 74 to 76. Here, for isolates BB12 and Cam_01, alternate alleles were incorrectly called as reference alleles using Nanopore data (Figure 3). For Cam_01, reference alignment bias due to low coverage may have contributed to the erroneous genotype call at CRT codon 74; BB12, however, had high overall coverage. Inspection of the alignments in this region revealed that for BB12, one of the resistance-associated variants gave rise to a homopolymer. No such homopolymer was introduced by the XHA mutation (Figure 3A), hence resulting in concordant haplotypes. Since Nanopore sequencing is subject to systematic error in homopolymeric stretches, this again demonstrates that caution should be exercised when potential homopolymeric tracts are encountered in loci selected for genotyping. However, we note that recent advances in Nanopore flow cell chemistry and improved basecalling algorithms will help mitigate these errors further.

**Figure 3:**
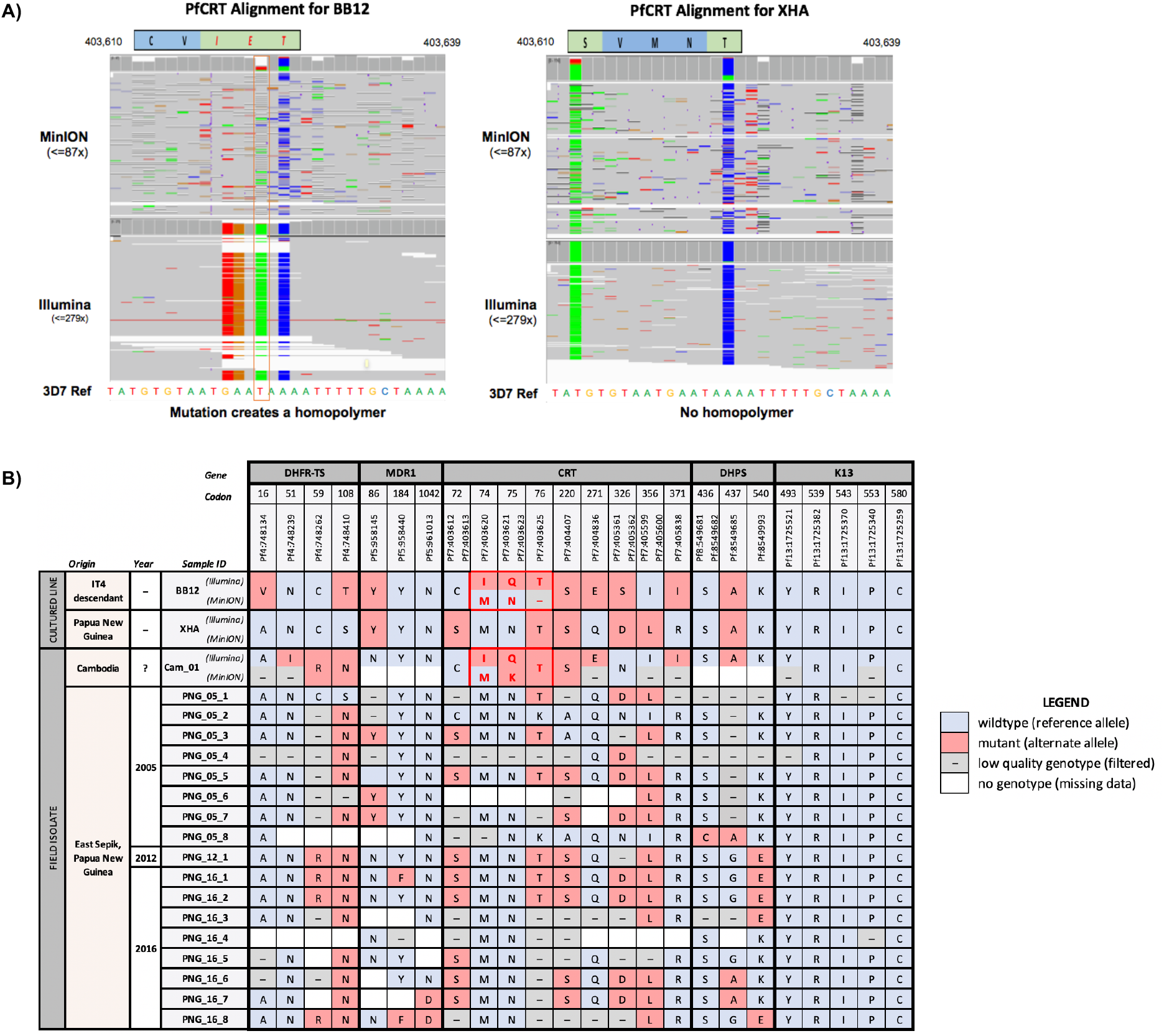
Drug resistance profiling of MinION field isolates. (A) Alignments of MinION sequence data across codons 72 to 76 of CRT for cultured lines BB12 and XHA. MinION and Illumina genotypes are discordant for BB12, where the mutation at position Pf3D7_07_v3:406325 has given rise to a homopolymeric tract. MinION and Illumina genotypes are concordant for XHA, for which there is no such homopolymeric stretch. (B) Drug resistance haplotypes for field isolates with breadth of coverage >65% at 1X. Isolates were genotyped at loci corresponding to key resistance-associated variants in genes crt, mdr1, dhfr, dhps and k13. Crt variants previously associated with resistance that were screened included T93S, H97Y, F145I, M343I and G353V however all isolates had wildtype alleles. Red boxes indicate resistance alleles, while wildtype alleles are indicated in blue. Missing genotypes (for which there was no sequence data) and low quality genotypes (which were removed after quality filtration) are shown in white and grey respectively.

Amino acid haplotypes spanning key resistance-associated variants across drug resistance-associated genes are presented in Figure 3B. Only Nanopore sequenced field isolates with a minimum breadth of coverage 65% at 1X are presented, and except for *kelch13*, only polymorphic loci within the dataset are shown. Some genotype calls were filtered due to low coverage. Missing genotype calls (occurring when no reads align to a particular locus) tend to be clustered, since genome-wide coverage for some Nanopore field isolates was uneven with no data captured for some genomic regions. These issues should be resolved as methods for parasite genomic DNA enrichment improve.

### Nanopore whole genome sequences are directly comparable to publically available P. falciparum Illumina data

Given that the majority of publically available *P. falciparum* genomes have been determined using Illumina short read sequencing, we asked whether genotypes obtained using Nanopore sequencing could be directly compared with these genotypes or whether Nanopore sequencing gave rise to platform-specific effects, thus preventing any biological insight from direct comparisons. We examined clustering patterns for Nanopore-sequenced field isolates (originating from PNG or Cambodia) relative to Illumina-sequenced isolates (originating from PNG, Cambodia, Vietnam, Laos or Thailand) from the MalariaGEN Pf3k/PNG dataset.

Each isolate was genotyped at a set of 742,365 high-quality SNVs identified thrugh an in-house reanalysis of the MalariaGEN P3k/PNG dataset. Genotype missingness filtration by isolate (<30%) and SNV loci (<10%) was performed to avoid possible biases introduced by imputation. Monomorphic loci were removed since they were uninformative, but all polymorphic SNVs were retained; SNVs were also not filtered by minor allele frequency to avoid ascertainment bias. After filtration, we retained 10 Nanopore- and 1040 Illumina-sequenced field isolates, genotyped at 51,421 polymorphic SNV loci.

Principal coordinates analysis (PCoA) was then performed on pairwise genetic distances between isolates, defined to be the proportion of successfully-genotyped loci with shared alleles for each pair of isolates. Projections of isolates onto two dimensions are visualised in Figure 4. Expected geographical clustering patterns emerged, with field isolates originating from PNG and South East Asia generally clustering distinctly. Nanopore- and Illumina-sequenced field isolates originating from PNG cluster together closely, suggesting that the different sequencing platforms do not impact the clustering analysis, with biological signals dominating instead.

**Figure 4.**
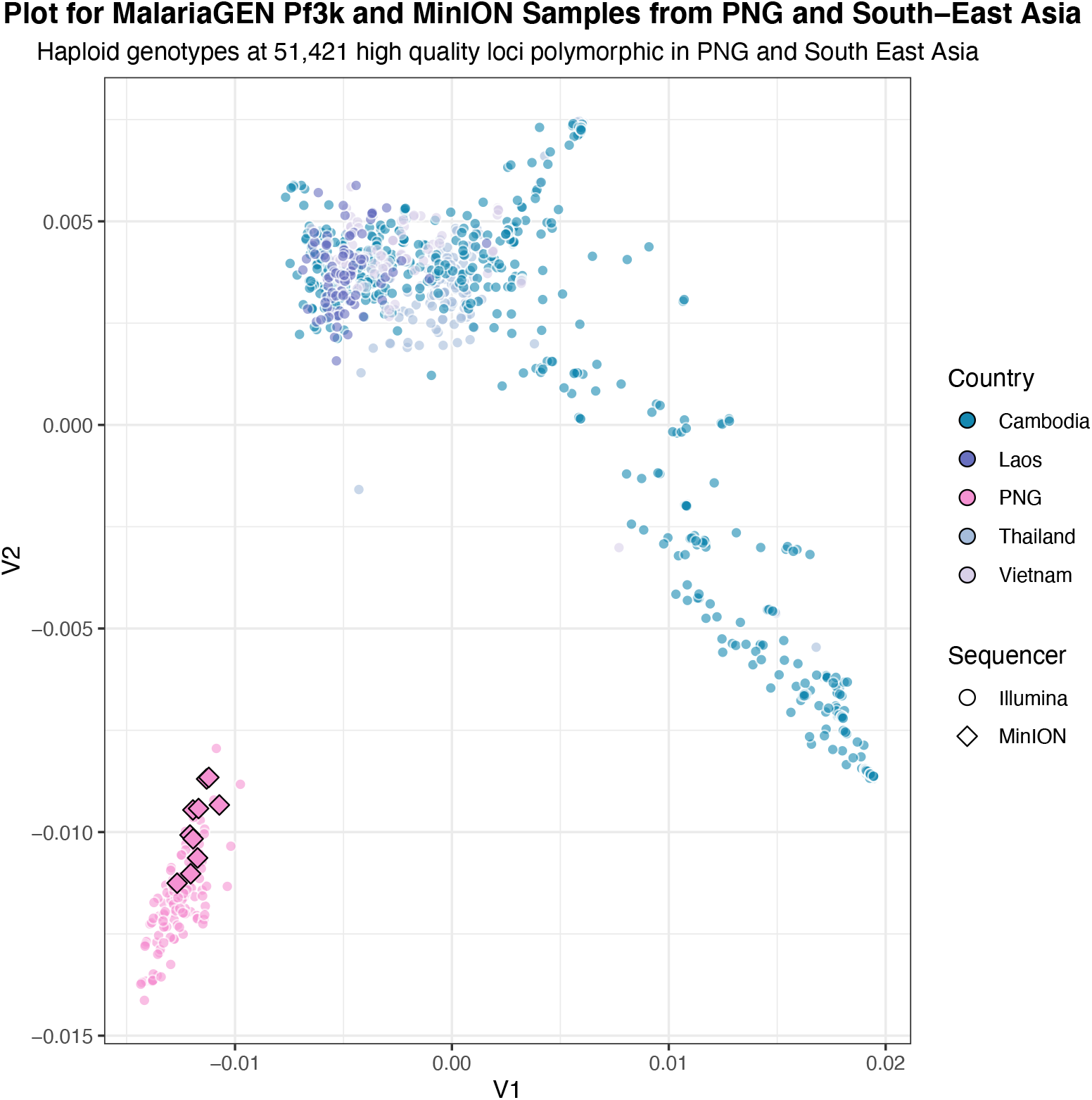
Clustering of Nanopore and Illumina-sequenced field isolate genomes from Papua New Guinea and South-East Asia (Cambodia, Laos, Thailand and Vietnam). Whole genome sequence data was used to genotype 742,365 high-quality SNVs obtained from an in-house reanalysis of the MalariaGEN Pf3k/PNG dataset. Ten Nanopore and 1,040 Illumina-sequenced field isolates were retained after samples with genotype missingness above 30% were filtered. Filtration of SNPs with missingness above 10% yielded 51,421 polymorphic SNVs. Pairwise distances between isolates, based on the proportion of the genotype shared between pairs, were calculated using the filtered set of SNVs. PCoA was performed on the resultant distance matrix to obtain a two-dimensional representation of the data. Each point represents a distinct isolate, with colours representing geographic origins, and shapes representing sequencing platforms. Nanopore- and Illumina-sequenced field isolates originating from PNG cluster together closely in the PCoA.

### De novo genome assembly using Nanopore long reads allows characterisation of highly polymorphic regions

*De novo* genome assembly can further improve characterisation of parasite genomes, enabling the inclusion of highly-polymorphic genomic regions that are typically blacklisted during variant calling from srWGS due to extreme polymorphism relative to the reference genome. The advent of lrWGS has greatly enchanced the continuity and completeness of *de novo* genome assembly, particularly in low-complexity and repetitive genomic regions that have been difficult to resolve with short read fragments (<1000bp)^30^.

To characterise repetitive and highly polymorphic genomic regions, we generated *de novo* genome assemblies using a combination of long and short read data. Long, continuous scaffolds were first generated using Nanopore long read data. Individual base errors, indels, block substitution events, gaps and local misassemblies in the scaffolds were then corrected using Illumina short read data, an approach that has also been used by others (https://github.com/nanoporetech/ont-assembly-polish). Summary statistics for the polished *de novo* assemblies are presented in Table 2.

**Table 2:**
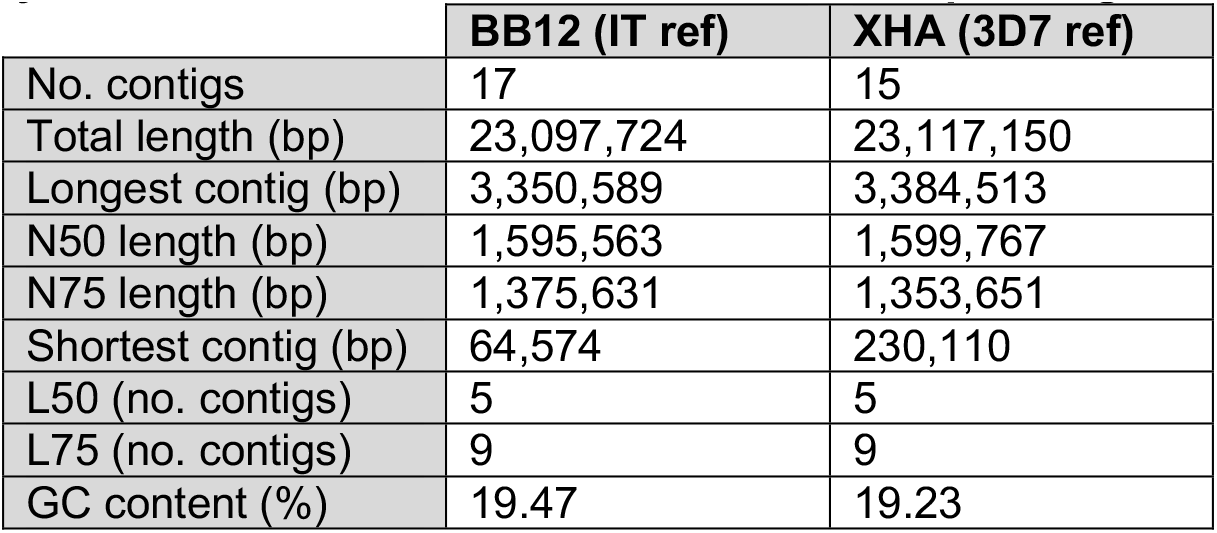
Summary statistics for *de novo* assemblies of *P. falciparum* genomes

The BB12 assembly was compared against the reference genome of its closest relative, the parent IT4 strain (version 4). However, for XHA, which is an isolate from PNG, no close relative reference genome is available so we compared the *de novo* assembly against the 3D7 reference genome; the 3D7 reference contains 14 complete nuclear chromosomes and has higher continuity than the IT4 reference genome. Assembly continuity for both BB12 and XHA was high, with all nuclear chromosomes except chromosome 5 spanned by a single contig (Figure S4). Breakpoints for chromosome 5, which was spanned by two contigs in both assemblies, varied for XHA and BB12. Alignment of the BB12 assembly against the IT4 reference genome (version 4) revealed that the two shortest contigs were spurious, multimapping to the subtelomeric regions of several nuclear chromosomes in addition to several contig fragments.

Although *de novo* contigs and reference chromosomes were generally concordant in core genomic regions, alignments in subtelomeric hypervariable regions were more fragmented. This fragmentation could reflect true variation between cultured lines and reference isolates (e.g. XHA vs 3D7), but may also be a marker of translocation and contig misassembly for homologous lines (e.g. BB12 vs IT4). A 150 kbp translocation from chromosome 13, for instance, is apparent in the downstream subtelomeric region of contig PfBB12_11.

The BB12 assembly allowed a more complete characterisation of chromosomes 6 and 13 relative to the IT4 reference genome (version 4). Since our BB12 assembly exhibited higher continuity than the IT4 assembly (version 4), we were able to localise fragment PfIT_00_11 to the subteolomeric region of chromosome 6, while fragments PfIT_00_4 and PfIT_00_10 were identified to be neighbouring subtelomeric flanks of chromosome PfIT_13 (Figure 5).

**Figure 5.**
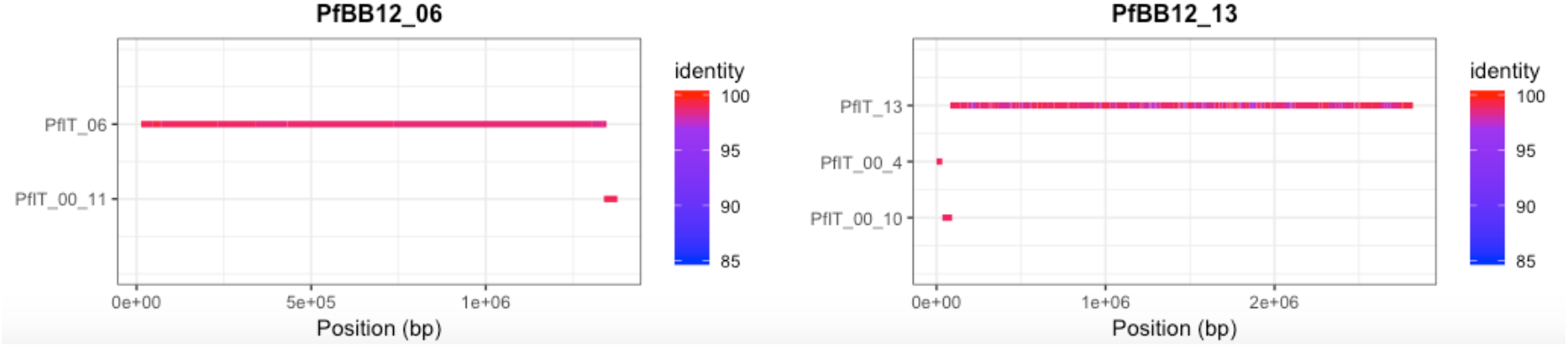
Enhanced continuity of MinION BB12 assembly, relative to the IT4 reference genome (version 4). More complete assembly of chromosomes 6 and 13 was enabled through long read sequencing. Fragment PfIT_00_11 has been localised to the subtelomeric region of PfIT_06, while fragments PfIT_00_4 and PfIT_00_10 have been identified as neighbouring subtelomeric regions of chromosome 12. One-to-one mappings between de novo contigs and reference genomes have been computed using nucmer (MUMmer, V.3.1^31^).

A genome-wide summary of annotated features in our assemblies for XHA and BB12 is shown in Table 3, with corresponding statistics for the 3D7 reference genome shown, as a comparison. The elevated number of pseudogenes identified relative to the 3D7 reference genome, in addition to the lower annotated gene density, suggests that open reading frames (ORFs) in the *de novo* assemblies were disrupted by mismatch and indel errors in a number of instances. Further, some annotated genes were fragmented, with multiple neighbouring annotations likely corresponding to the same gene. However, we were still able to characterise a large number of genomic features in our assemblies.

**Table 3:**
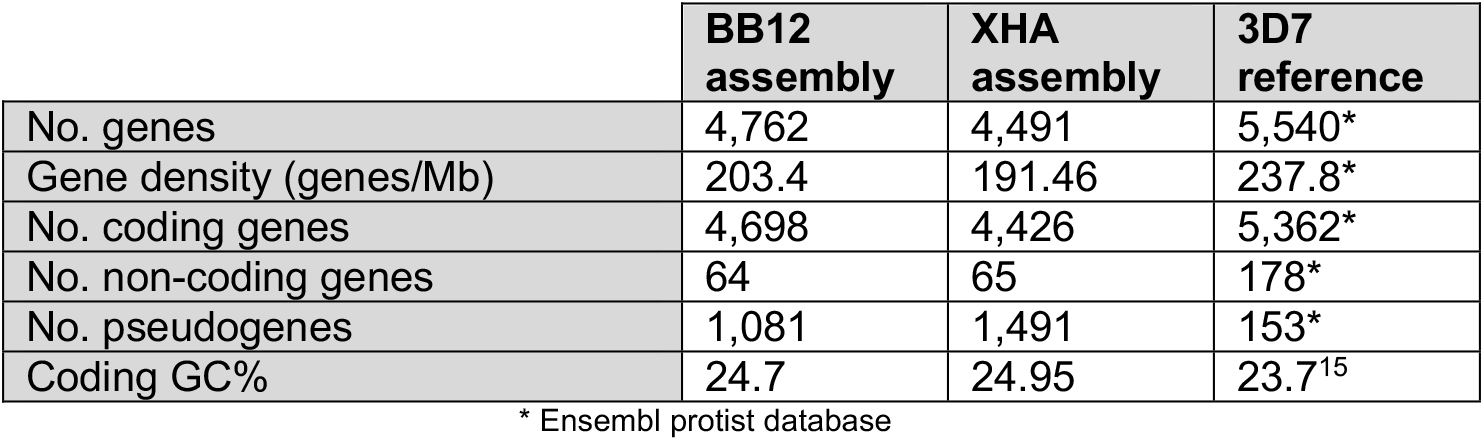
Summary of annotated genomic features for *de novo* assemblies of *P. falciparum* genomes compared to the 3D7 reference genome.

We identified 27 complete, 16 split and 11 partial *var* genes for BB12, and 18 complete, 16 split and 15 partial *var* genes for XHA. Domain structures for all complete and partial *var* genes are shown in Figure S5.

### Improved structural variant calling using long read data

Approaches utilising srWGS tend to provide limited resolution for the detection of structural variants (SVs)^32^. LrWGS has emerged as a promising tool for SV detection, particularly complex SV. Concordance rates across three trialled SV pipelines are presented in Figures 6A and 6B, focussing on SVs longer than 200bp to avoid systematic errors associated with Nanopore sequencing^33^. SVs detected using long reads only (Sniffles^33^) and our *de novo* assemblies (Assemblytics^34^) exhibited reasonable levels of concordance, with approximately 30% of high-quality SVs consistent across both pipelines. SVs detected using short reads only (GRIDSS^35^) had very little concordance with either Sniffles or Assemblytics called SVs. Short read SV calling (GRIDSS) identified fewer high-quality SVs longer than 200 bp, with the vast majority of high-quality SVs within the length range 30-200 bp. Longer SVs failed to pass default quality filtration parameters. Distributions of SV types and lengths for the three pipelines are shown in Figures 6C and 6D.

**Figure 6.**
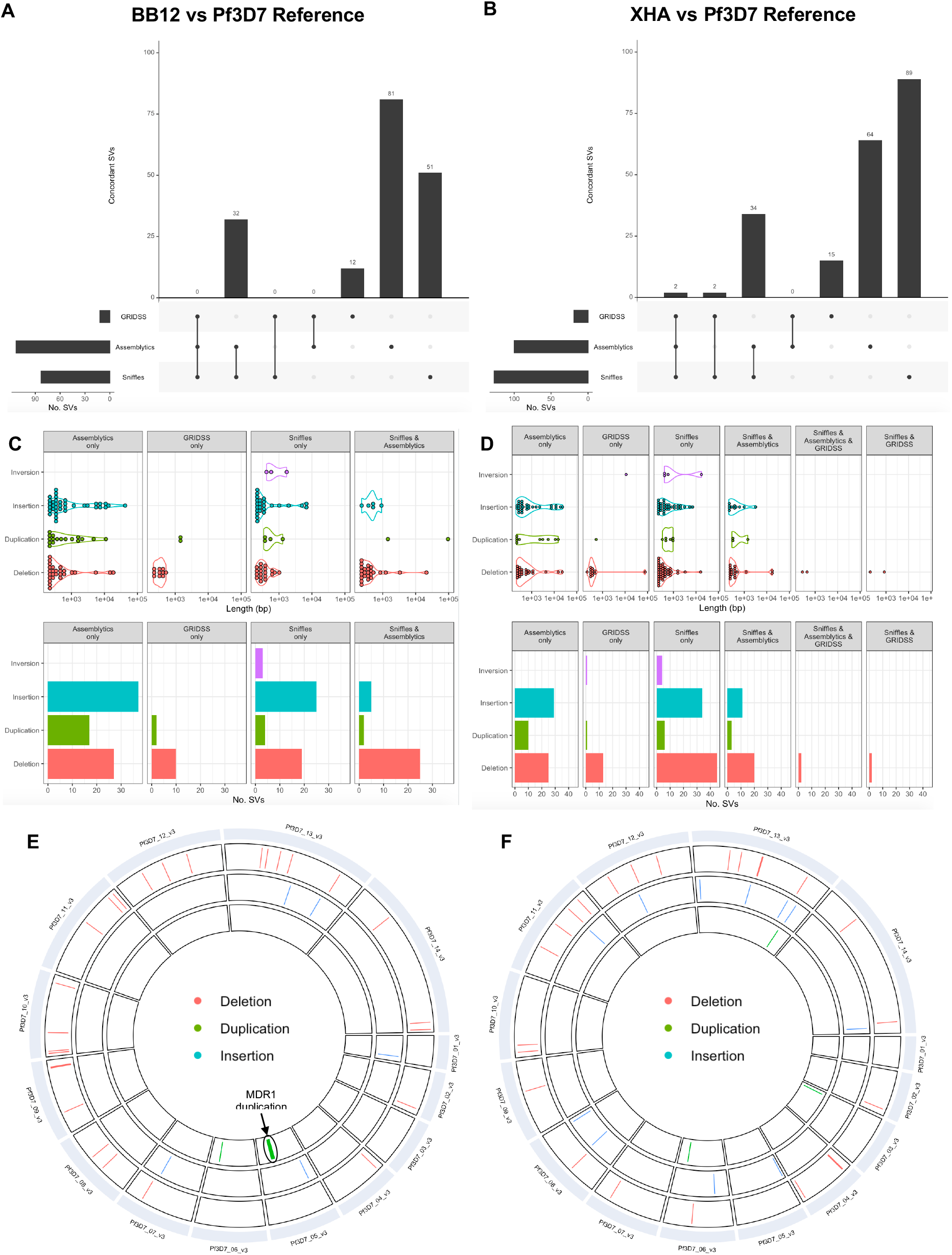
Comparison of structural variant calling piplines using long reads only (Sniffles), short reads only (GRIDSS) and our de novo assemblies (Assemblytics). Overlaps in the sets of high-quality SVs identified by each pipeline are BB12 and XHA are shown in (A) and (B) respectively. SVs identified by different pipelines are considered equivalent if they overlap by at least 1bp, correspond to the same SV class and have similar lengths (i.e. the length of the shorter SV is at least 75% of that of the longer SV) (Text S8). Sniffles (which uses long reads only) and Assemblytics (which is informed by our de novo assemblies) show reasonably high concordance, while GRIDSS (which uses short reads only) exhibits very little concordance with either Sniffles or Assemblytics. SV types and lengths for BB12 and XHA are shown in (C) and (D) respectively. SV length and type distributions are generally quite similar for Sniffles and Assemblytics. Genome-wide SV maps, restricted to SVs concordant across Sniffles and Assemblytics, are shown for isolates BB12 (E) and XHA (F) respectively. Both Sniffles and Assembytics detected the known MDR1 duplication in BB12.

We then screened high-quality SVs to identify functionally-relevant variation. For BB12, a 96kbp duplication on chromosome 5 spanning *mdr1* (PF3D7_0523000) was identified by both Sniffles and Assemblytics, but not GRIDSS. A quantitative real-time PCR (qPCR) assay confirmed the presence of the duplication spanning *mdr1* (data not shown). The amplification of *mdr1* has been associated with resistance to a range of antimalarial drugs, including artemisinin, lumenfantrine, mefloquine and halofantrine^36^. While this SV was successfully captured by lrWGS data, srWGS failed to identify this amplification. No gene amplifications were observed for other drug resistance markers.

The results demonstrate enhanced sensitivity and resolution of SV calling pipelines leveraging lrWGS data, compared to approaches using srWGS data in isolation. Genome-wide SV maps for isolates BB12 and XHA, constructed using SVs concordant across Sniffles^33^ (based on lrWGS mapping to a reference genome) and Assemblytics^34^ (based on *de novo* assemblies) are shown in Figures 6E and 6F respectively.

## DISCUSSION

The Oxford Nanopore MinION sequencer is an attractive platform for sequencing malaria parasites in the field and laboratory. However it is known to have higher error rates in comparison to Illumina platforms. In addition, *P. falciparum* has an AT rich genome, with a higher probablility of homopolymers, known to be the cause of high error rates in Nanopore WGS. Here we benchmarked Nanopore long reads against Illumina short reads to reveal the utility of Nanopore for WGS of *P. falciparum* laboratory and field isolates. The results reveal a smaller baseline error rate than expected for haploid SNP genotyping using Nanopore long read sequencing. Discordant allele calls between Nanopore and Illumina data were more frequent in homopolymer regions greater than 6 bp, which are known to have poor sequence quality using the R9 (R9.4 and R9.5) Nanopore flow cells used here, and whose occurrence is expected to decrease with new Nanopore pore designs, as this is an active area of development. We optimised an enrichment and library preparation for whole blood-derived DNA samples enriched for parasite DNA after treatment with a *McrBC* digestion and rWGA, which could also be used for other *Plasmodium spp*. such as *P. vivax* and even other pathogens. Through sequencing field samples with a range of infection densities, we provide recommendations for selecting *P. falciparum* isolates for Nanopore WGS. While early runs resulted in outputs of 2-3 Gb, at the time of writing, we are generating up to 18 Gb of data per run using the more recently available SQK-LSK109 kit (unpublished data), which would increase coverage by almost ten fold and therefore allow multiplexing of up to 12 samples. In addition, we report streamlined analytical pipelines for SNP genotyping and de novo assembly, which yielded complete chromosomes including highly polymorphic subtelomeric regions and large SVs.

Removing dominant sources of variation (namely, indels; subtelomeric polymorphisms and heterozygosity) significantly increases the accuracy of genotyping with the Nanopore platform. The low error rates (<1% off filtered calls) computed were based on haploid SNP genotyping and are significantly lower than reported error rates for diploid or ‘double haploid’ SNP genotyping, when heterozygous calls are taken into account^29^. In addition, transition-type errors (G to A or C to T), were more frequent than transversion-type errors, possibly reflecting errors in basecalling. While the accuracy of Nanopore reference calls was high, alternate allele calls were found to be less reliable at low depths of coverage; however, overall discordance rates remained low since reference calls far outnumbered alternate calls. Reliably detecting alternate alleles at low depths of coverage may be problematic for Nanopore sequencing in some applications.

The results demonstrate the utility of the Nanopore MinION platform as a tool for drugresistance profiling, even at very low depths of coverage (breadth of coverage ≥65% at 1X), granted caution is exercised around potential homopolymeric tracts flanking target loci where error rates were higher. High concordance with Illumina data was observed except for *crt* codons 72-76, which contains the chloroquine resistance haplotype. XHA (SVMNT) showed high concordance between Nanopore and Illumina data, but there were several discordant positions for BB12 (CVIET), due to a T to A mutation within codon 75 that creates a homopolymer. As these haplotypes are more common in some parts of the world, caution is warranted when genotyping *crt* using Nanopore sequencing. Some drug resistance genes and samples were genotyped more consistently, e.g. *k13* was fully genotyped in almost all samples, whilst *crt* had more missing data due to the introduced homopolymer. With increasing information on the variant frequency in target populations however, missing genotypes can be imputed^37^ and flow cell chemistry has improved to allow more accurate sequencing of homopolymers. Despite missing data for some samples, a large number of PNG isolates from different time points were successfully genotyped for all genes. We observed a high prevalence of resistance markers for chloroquine and antifolate drugs^38, 39^ yet a complete lack of *kelch13* mutations associated with artemisinin resistance. This is critical information for PNG given the recent reports of a small number of C580Y mutant parasites in Wewak in 2016^40^, only 50 km from the collection site for the samples sequenced here. The additional ability for Nanopore to return results within 48 hrs at field sites, rather than weeks or months after samples have often left the country of collection means that field relevant results can be returned in a time frame relevant for decision making regarding local treatment regimens or programmatic interventions.

As platform technologies change and improve, it will be important to ensure that new data can be directly compared to other, previously generated and public datasets^24^, in order to track parasite evolution in response to changes in transmission and different selection pressures. While our analysis of discordance rates between Nanopore MinION and Illumina (haploid) SNP genotyping suggests the presence of some systematic differences between the two sequencing platforms, these effects may not be of a sufficient magnitude to significantly skew population genomics analyses. Our results are a preliminary validation, suggesting the potential viability of population genomics analyses combining Nanopore and Illumina data.

We also demonstrate the ability of Nanopore long read sequencing data to resolve hypervariable and low complexity genomic regions that have been difficult to characterise with short reads. Nanopore data allowed the construction of highly continuous genomic scaffolds that could then be polished with Illimina data to correct small-scale errors, including indels, individual base errors, block substitution events and local misassemblies. We generated highly complete and continuous *de novo* assemblies, validated through a comparison of BB12 against its ancestral reference strain IT4^19, 20, 41^. Although a number of *de novo* assemblies for *P. falciparum* drawing on single-molecule real-time (SMRT) long read sequencing using the PacBIO platform have been published^41^, we presented the first *de novo* genome assemblies for *Plasmodium falciparum* generated using Nanopore sequence data.

While structural variants (SVs) play an important role in the genomic diversity of *P. falciparum*^42^, shortcomings persist in the detection of SVs with short-read sequencing^32^. Long-read sequencing has the potential to enhance the sensitivity and specificity of SV detection, however, applications of long-read sequencing to characterise structural variation in the *P. falciparum* genome^43^ have been limited. Studies have focused on comparisons between reference genomes and *de novo* genome assemblies generated from PacBIO SMRT long-read sequencing data. We demonstrated the enhanced ability of Nanopore long reads to capture SVs relative to short reads alone by comparing pipelines for SV detection relying on Illumina only mapped to a reference genome^35^; Nanopore only mapped to a reference genome^33^, as well as *de novo* genome assemblies^34^. Of note, we detected a 96kbp duplication on chromosome 5 spanning *mdr1* in BB12, that was not detected by short read sequencing data. The ampliciation of *mdr1*, which is associated with multidrug resistance^36^, was confirmed by qPCR. While further research is required to generate robust sensitivity and specificity estimates for SV detection, Nanopore sequencing is a promising tool for fine-scale mapping of complex SVs in the *Plasmodium falciparum* genome.

Nanopore long read sequencing using the MinION portable device has been proposed as a useful platform for real time portable sequencing solutions, with applications to malaria surveillance. Here we have optimised Nanopore sequencing protocols and data analysis for high quality *P. falciparum* WGS. The results described offer insight into the quality and utility of Nanopore long read sequencing, and its inherent limitations. Following filtering of low confidence variation we observed acceptably low error rates and very high concordance of resulting genome wide (haploid) SNP genotypes for both pure *P. falciparum* DNA and field isolates contaminated with high amounts of human DNA. Long reads generated using this platform improve whole genome assembly and the detection of hypervariable regions and SVs. Nanopore genomes are directly comparable to publically available Illumina genomes and reveal novel insights of practical importance for malaria control programs including population structure and drug resistance. Finally, the Nanopore sequencing platforms also offers the potential for improving research equality in the many countries still battling malaria by enabling sequencing in country, giving research power to local malaria researchers and national malaria control programs to inform programmatic decisions.

## METHODS AND MATERIALS

### Parasite isolates

Pure *Plasmodium falciparum* genomic DNA was obtained from three culture-adapted isolates including the reference strain 3D7; BB12, a descendant of the IT strain^19, 20^, and XHA, a culture-adapted isolate from PNG^18^. Annotated genome assemblies are available for 3D7^44^ and IT^41^. *P. falciparum*-infected blood samples (n=33) were collected in cross-sectional surveys in Papua New Guinea, where malaria transmission and parasite genetic diversity is high^45^ and in Cambodia where transmission is low^46^ with high levels of multidrug resistance^47^. The research was approved by the PNG Institute of Medical Research Institutional Review Board No. 1116, Medical Research Advisory Council of PNG No. 1121, Cambodian National Committee on Health Research No. 265 NECHR and Walter and Eliza Hall Institute of Medical Research Human Research Ethics Committee Nos. 1303 and 1504.

### DNA extraction, parasite DNA enrichment and library preparation

For cultured *P. falciparum* lines, DNA extraction was performed using DNeasy Blood & Tissue Kit (Qiagen, Germany) according to the manufacturer’s instructions. Extracted DNA was purified and concentrated as described previously^48^, with SPRIselect beads (Beckman Coulter, Australia) used in lieu of Ampure XP beads. Briefly, 0.45 volume re-suspended beads were added to 1 volume of sample and incubated at room temperature for 10 minutes. After incubation, the bead/DNA mixture was placed on a magnetic rack for 5 minutes. The supernatant was removed without disrupting the DNA-bound beads, and the beads were then washed twice with 200 μl of 80% ethanol. After incubating from an additional 10 minutes at room temperature, the bound DNA was eluted with 60 μl of 10 mM Tris buffer. The quantity and quality of extracted DNA were measured using a Qubit Fluorometer (Invitrogen Life Technologies, USA) and Nanodrop2000 (Thermofisher, USA) respectively. Size distributions were then analysed using Tapestation (Agilent Technologies, USA). Based on these results, 0.2pmol of DNA was processed with the 1D genomic sequencing kit SQK-LSK108 following the manufacturer’s instructions (ONT, UK).

To optimise library preparation for field samples, mock infections were prepared to mimic 1% parasitemia by spiking 2 ng (0.083ng/ μL) of 3D7 parasite DNA with 73 ng (3.066/ μL) of commercial human DNA in a total volume of 24 μL. This is approximately 6.35 parasites/human genome and equivalent to a 1% parasitaemia. Both methylation-dependent digested (*McrBC*) and undigested mock infections were tested to verify the efficacy of the enrichment protocol (described in detail below). Library preparation was done using the 1D genomic sequencing kit following the manufacturer’s instructions (Cat # SQK-LSK108, ONT, UK), however, we assessed various adjustments to the post-whole genome amplification (WGA, see below) and end prep purification. Following WGA, we trialled two methods of purification: 2.5V ethanol precipitation and 1.8V SPRI beads. Following end prep, we further tested 0.45V and 1V volumes of bead clean-up. The optimal protocol was found to involve; 1.8V SPRI bead purification after WGA and 0.45V bead purification following end prep. All field samples were subject to this protocol. Three to four field samples were multiplexed for each run by indexing with the native barcoding kit (Cat # EXP NBD103, ONT, UK).

For field isolates, DNA was extracted from dried blood spots using the FavorPrep™ 96-Well Genomic DNA Extraction Kit (Favorgen, Taiwan). In order to reduce costs and minimise the amount of human DNA contamination in the sequencing output, we developed a novel potentially universal enrichment strategy that is dependent on low methylation of the *Plasmodium* genome (less than 1%)^49, 50^, relative to the high levels of methylation in the human genome (60-90%)^51^. The protocol involves 2-steps: a restriction digest that targets methylated cytosines followed by whole genome amplification (WGA) using the Phi DNA polymerase primed with random hexamers. Briefly, 6 μL of DNA was subject to restriction digestion with *McrBC* (New England Biolabs, US) at 37°C for 1 hour, and halted by incubation at 80 °C for 20 minutes. Gel electrophoresis confirmed a smear for human DNA alone, indicating that human DNA had been digested, and maintenance of high molecular weight parasite DNA. Then, 2 μL of each field sample was subject to whole genome amplification using the illustra Genomiphi V2 DNA amplification kit (GE Healthcare Life Sciences, Australia) for 2 hours at 30°C followed by heat inactivation at 65°C for 10 minutes. Amplified DNA samples were then purified using 1.8V bead purification (Beckman Coulter, Australia) and further treated with T7 Endonuclease I to remove branch structures produced during whole genome amplification. All samples were amplified in duplicate-triplicate and pooled prior to library preparation.

To assess the efficacy of enrichment for mock and natural infections, we performed duplex qPCR, targeting the *P. falciparum 18S rRNA* genes^52^ and the human *Plat1* gene^53^. We combined 0.2 μL (150 nM) of both forward and reverse primers; 0.45 μL (350 nM) of Taqman probes; 6.5μL of Taqman Fast Advanced Mastermix (Applied Biosystem, USA) and 4 μL of target DNA, comprising of undiluted original DNA or a 1:100 dilution of the amplified product. Details of the relevant primers and probes are presented in Table S1. To quantify parasite and human copy numbers, standard curves were generated by preparing 10 fold dilution series of plasmid DNA containing parasite *18SrRNA* gene^54^ and from human genomic DNA (Cat No. G304, Promega, Australia). Thermal cycling was performed on a Light Cycler 480 II (Roche, Switzerland). The reaction volume was heated to 95° for 15 minutes, followed by 40 cycles of 95° for 15 seconds and 60° for 1 minute.

### Nanopore sequencing and raw data analysis

Sequencing runs, each spanning 48 hours, were performed using Nanopore flowcells MIN106/MIN107 (R9.4 and R9.5) and MinKNOW software V1.7.10-1.11.5 (ONT, UK). Basecalling and demultiplexing were performed using the ONT Albacore (V1 for isolate XHA and V2.1 for others) to obtain reads in *fastq* format. Adapters were trimmed from basecalled reads using *Porechop V0.2.1*^55^ with default parameters. Read length and quality summaries were subsequently generated with *NanoPlot V0.16.4*^56^. Trimmed reads were then aligned to the *P. falciparum* 3D7 (version 3)^44^ reference genome as well as a concatenated *P. falciparum* 3D7, human HG19 and *P. vivax* P01 reference^57^, using *Minimap2 V2.4*^58^ with the *map-ont* preset. *Samtools V1.7*^59^ utilities were used to sort and index and resultant alignments to obtain sorted bam files. Coverage statistics and the proportions of parasite and human DNA were determined using *samtools depth* and several in-house helper scripts. Locus-wise alignment summaries were generated using *samtools mpileup* V1.7^59^, using a base quality threshold of 7. Alignment summaries were then used to call haploid variants using the *bcftools multiallelic-caller* V1.8^60^, in both genotyping and discovery modes. Indels were removed using *vcftools V0.1.13*^61^ and functional annotation of the resultant SNPs was performed using SnpEff V4.1l^62^. Quality filtration based on the depth of coverage at each locus and the proportion of reads supporting the called allele was performed using an in-house script (Text S1). A schematic of this pipeline for processing Nanopore data is shown in Figure S1A.

Statistical analysis to determine associations between sample parameters and Nanopore sequencing output was done using the R package *ggpubr*^63^. Linear regression lines, with 95% confidence intervals, were generated with the function *ggscatter*, and Pearson’s correlation coefficient was computed with the *stat_cor* utility.

### Illumina sequencing and raw data analysis

We also performed Illumina sequencing of cultured *P. falciparum* isolates 3D7 (unknown origin), BB12 (Brazil) and XHA (PNG) which contain only *Plasmodium* spp. DNA, in addition to three Cambodian field isolates which are contaminated with human DNA. These samples were used to benchmark MinION sequencing against the widely used Illumina sequencing approach. Library preparation for field isolates required a preliminary parasite DNA enrichment step (see above). Sequencing was performed as per the Truseq Nano DNA sample preparation protocol (Illumina Inc., USA). Briefly, 200 ng of DNA was subject to shearing, endrepair, A-tailing and adapter ligation, followed by enrichment with 15 cycles of PCR using BIORAD T100 Thermal Cycler (Australia). The mean insert size was analysed using Tapestation (Agilent Technologies, USA) with D1000 Screen Tape (Cat No. 5067-5582). Libraries were then sequenced using the Nextseq 500 platform (Illumina Inc., USA) generating 75 bp paired-end reads with 6-base index read. Data analysis was done using an in-house pipeline (http://github.com/bahlolab/pf_variant_calling_pipeline) in accordance with GATK best practices^64^ (Text S2).

### Data analysis

#### Baseline error rates

To quantify false discovery rates for *de novo* SNP characterisation, variant calling was performed in discovery mode for both Nanopore and Illumina data to detect novel (haploid) SNPs in our cultured 3D7 isolate, relative to the 3D7 reference genome. Sequencing data for both Nanopore and Illumina comprised of mock infections. For Nanopore, we retained SNPs with minimum depth of coverage 10X and at least 75% of reads supporting the called allele. Illumina SNPs were filtered in accordance with GATK best practices (that is, QD>=2, MQ>=40, FS<=60, SOR<=3, MQRankSum>=−12.5 and ReadPosRankSum>=−8). To avoid detecting true variation between our cultured line and the isogenic reference strain that had arisen due to mitotic recombination *in vitro*^22^, we considered only coding SNVs in essential genes that would have been unlikely to differ between the cultured and reference strains (Text S3).

#### SNV calling

Validation of SNV genotyping was done by first comparing *P. falciparum* isolates sequenced in-house using both Nanopore and Illumina platforms. Variants were called at 742,365 validated SNV loci obtained from an in-house reanalysis of the MalariaGEN Pf3k dataset (release 5), pooled with an additional set of 149 field isolates from Papua New Guinea^25^. Discordance rates between Nanopore and Illumina genotypes for cultured lines 3D7, BB12 and XHA, and the two Cambodian field isolates were quantified using an in-house script. Error rates in homopolymeric stretches more than 6bp in length, identified using GATK VariantAnnotator (V3.5) were compared against non-homopolymeric stretches. Filtration parameters for haploid Nanopore SNV genotyping (depth of coverage at least 2X, 75% of reads supporting the called allele) were subsequently deduced.

Drug resistance profiling was then performed for field isolates with at least 1X coverage in 65% of the genome. Genotypes were then screened for known resistance-associated variants in multiple genes including *crt* (PF3D7_0709000), *mdr1* (PF3D7_0523000), *dhfr* (PF3D7_0417200), *dhps* (PF3D7_0810800) *and kelch13* (PF3D7_1344700).

To determine whether there were any platform-specific effects in the SNV genotypes called from WGS data, we then performed a principal coordinates analysis (PCoA) of in-house Nanopore field isolate data (PNG) and Illumina field isolate data obtained from the MalariaGEN Pf3k/PNG dataset (Asia Pacific including PNG and Cambodia) based on 51,421 high-quality polymorphic SNVs that were present in those countries (Text S4).

#### De novo genome assembly

Raw Nanopore reads for BB12 and XHA were first assembled into a scaffold using *Canu V.1.3*^65^ using default parameters. Five iterations of consensus polishing were then performed with *Racon V.0.5.1*^66^, which used the mapping of uncorrected reads to the scaffold, computed using *minmap2 V2.4*^58^ to generate consensus sequences. Draft assemblies, constructed from Nanopore data only, were further polished with Illumina sequencing data. Illumina reads processed with *Trim Galore V.0.4.4*^67^ were first mapped against draft assemblies using *bwa mem V.0.7.13*^68^. Individual base errors, indels, block substitution events, gaps, local misassemblies and ambiguous bases in the draft assemblies were then corrected using *Pilon V.1.22*^69^ to obtain hybrid Nanopore-Illumina assemblies. A schematic of this pipeline is shown in Figure S1B. Draft assemblies for BB12 were benchmarked against its ancestral reference strain IT (Text S5).

Hybrid assemblies were compared against *P. falciparum* reference genomes using *nucmer* (*MUMmer, V.3.1*^31^), configured to identify one-to-one mappings between de novo contigs and reference chromosomes. Gene annotation was then performed with the *Companion* pipeline (July 2019 web server version)^70^. Domain classification of annotated *var* gene candidates was performed using the *VarDom 1.0 Server*^71^ (Text S6).

#### Structural variant (SV) calling

To assess the utility of Nanopore lrWGS for the characterisation of SVs in *P. falciparum*, we performed a comparison of three distinct SV-calling pipelines for cultured lines BB12 and XHA: (i) Short read SV-calling with GRIDSS^35^, which combines split-read, read-pair and assembly approaches, (ii) long read SV-calling with Sniffles^33^ and NGLMR^33^, which employs a split-read approach, and (iii) SV detection through direct comparison of our *de novo* assemblies and a reference genome using Assemblytics^34^ (see Text S7 for details). The rationale for using GRIDSS as a benchmark for short read SV calling was two-fold: in addition to combining a number of common approaches to short read SV calling, GRIDSS was found to provide high precision across a range of SV types in a recent evaluation of over 60 short read SV detection algorithms^72^. Concordance rates across the three methodologies were then computed (Text S7). Systematic indel errors in Nanopore basecalling can lead to an overrepresentation of small deletions in low complexity and homopolymers, hence, we considered only SVs with length at least 200bp, in line with general recommendations for Sniffles^33^. Several spuriously large SVs, spanning almost entire chromosomes in some cases, were detected by both GRIDSS and Sniffles. Hence, only SVs with length below 300,000 bp were retained.

## Acknowledgements

We are grateful to the volunteer communities and field teams and laboratory staff of PNG Institute of Medical Research and Institut Pasteur Cambodia for their involvement in sample collections. Thank you also to Celine Barnadas for facilitating access to Cambodian samples and Stuart Lee for assistance with genomic analyses.

## Funding

This research was funded by grants from the Australian National Health and Medical Research Council (NHMRC) GNT1163420 and Department of Foreign Affairs and Trade. Samples from PNG were collected with funding from the NIH NIAID International Centres of Excellence in Malaria Research (ICEMR) South West Pacific (U19 AI089686) and a Bill & Melinda Gates Foundation TransEPI grant. LJR, IM and MB are supported by NHMRC Research Fellowships (GNT1161627, GNT1155075, GNT1102971). DLG is supported by the NIH NIAD International Centres of Excellence in Malaria Research (ICEMR) Asia Pacific (U19 AI129392-01) and the NHMRC Australian Centre of Research Excellence in Malaria Elimination Grant 1134989. The authors acknowledge the Victorian State Government Operational Infrastructure Support and Australian Government NHMRC Independent Research Institute Infrastructure Support Scheme (IRIISS).

## SUPPORTING MATERIALS

Razook and Mehra *et al*. Real time, field-deployable whole genome sequencing of malaria parasites using nanopore technology

## SUPPORTING METHODS

**Text S1.** Details of Nanopore SNV quality filtration

**Text S2.** Pipeline for processing Illumina data

**Text S3.** Essential genes for baseline error rate analysis

**Text S4.** Principal coordinates analysis (PCoA) of isolates from PNG and South East Asia

**Text S5.** Benchmarking of BB12 *de novo* assemblies against the IT reference genome

**Text S6.** Gene annotation and *var* gene domain classification

**Text S7.** Structural variant calling pipelines

### Text S1: Details of Nanopore SNV Quality Filtration

MinION genotype calls were assessed individually for each sample using two locus-wise quality metrics:

- The number of high-quality bases mapping to the locus (CovDepth), and
- The proportion of high-quality reads supporting the called allele (PropReads).

Both of these metrics were extracted from the DP4 tag, as described in the VCF file specification, which encodes the breakdown of high-quality bases mapping to the forward/reverse reference/non-reference alleles:

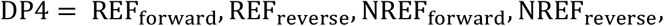

where (N)REF_forward_ and (N)REF_reverse_ denote the number of high-quality bases mapping to the forward and reverse (non-)reference alleles respectively. The DP4 was parsed using string manipulation functions in GNU *awk*, and quality metrics were calculated using the following formulae:

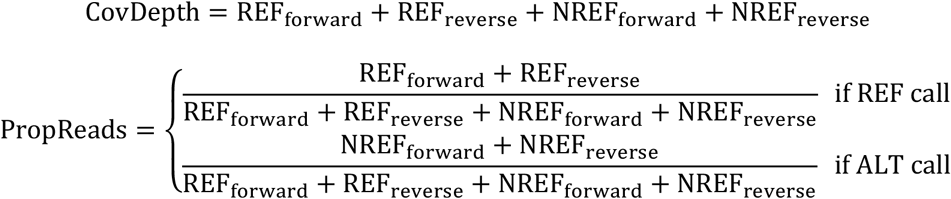

Since the DP4 tag considers only high-quality bases, CovDepth was lower than depth of coverage encoded in the DP tag for some loci.

### Text S2: Pipeline for Processing Illumina Data

Isolates sequenced in-house on the Illumina platform were analysed in accordance with GATK best practices. Briefly, Illumina adapters were identified using *Picard Tools MarkIlluminaAdapters V2.17.3*^1^ and reads were aligned against the P. *falciparum* 3D7 (version 3) reference genome using *bwa mem V0.7.13*^2^. Indels were realigned using the *GATK IndelRealigner V3.5.0*^3^, and duplicate reads were identified with *PicardTools MarkDuplicates*. Base quality score recalibration was the performed using the *GATK BaseRecalibrator*. Haploid variants were called using the GATK *HaplotypeCaller* in both DISCOVERY mode (which outputted all detected variants) and GENOTYPE_GIVEN_ALLELES mode (which genotyped isolates at a specific set of loci).

### Text S3: Essential Genes for Baseline Error Rate Analysis

Essentiality of genes for *in vitro* asexual blood stage development can be quantified using the piggyBac insertion mutagenesis score (MIS), which ranges from 0 for essential genes to 1 for dispensible genes^4^. To quantify false discovery rates for Nanopore and Illumina SNP calling, we restricted our attention to novel coding variants detected in the top quantile of essential genes (n=1356), with MIS in the range 0 to 0.142. Coding effects of each SNP were predicted using the R package *VariantAnnotation* (V1.32.0)^5^.

### Text S4: Principal Coordinates Analysis (PCoA) of Isolates from PNG and South-East Asia

Nanopore-sequenced field isolates were genotyped at 742,365 high-quality SNP loci obtained through our in-house reanalysis of the MalariaGEN Pf3k/PNG dataset. Only Nanopore genotypes with depth of coverage at least 2X and a minimum of 75% of reads supporting the called genotype were retained. Single-sample VCF files for Nanopore-sequenced field isolates were merged into a multisample VCF file using *vcftools V0.1.13*^6^. Multisample VCF files for both the MalariaGEN Pf3k/PNG datasets and our Nanopore-sequenced field isolates were then converted into *gds* format^7^ and imported into R statistical software^8^.

Isolates with genotype missingness rates exceeding 30% were removed, yielding 10 MinION-sequenced field isolates (all originating from Papua New Guinea) and 1040 Illumina-sequenced field isolates. Variant-level filtration was then performed to retain only SNP loci with missingness <10%, leaving 298,112 variants. SNPs monomorphic across the filtered panel of field isolates were removed to yield 51,421 polymorphic high-quality SNPs. These filtration steps were performed using *SeqArray V1.22.3*^9^.

Pairwise distances between the resulting genotypes, defined to be the proportion of differing SNP sites between each pair of isolates, were computed using the *dist-gene* function in the R package *ape V5.2*^10^; missing sites were excluded in pairwise comparisons. Principal coordinates analysis (PCoA) was performed on the resultant pairwise distance matrix with the base R function *cmdscale* (V3.6.2) with default parameters to examine clustering patterns stratified by geographic origin and sequencing platform.

### Text S5: Benchmarking of BB12 *de novo* Assemblies Against the IT Reference Genome

Since the cultured line BB12 is a descendent of the IT strain of *Plasmodium falciparum*, we expect the core nuclear genomes of BB12 and IT to exhibit a high degree of similarity. Our preliminary benchmark for *de novo* assembly was thus a contig-level alignment of a potential BB12 assembly to the IT4 reference genome (version 4)^11^. We sought to minimise indel and mismatch rates for draft BB12 assemblies compared (Figure S1B) to the IT4 reference genome, generating quality summaries for various pipeline stages using *QUAST V.5.0.2*^12^ (web server, May 2018 version).

The initial scaffold, generated using *Canu V.1.3*^13^ from raw Nanopore reads, had high continuity, with all but one nuclear chromosome spanned by a single contig and two spurious contigs; however, mapping between the scaffold and the IT4 reference genome was poor (Table M1). Five iterations of consensus polishing with *Racon V0.5.1*^14^ substantially improved alignment against the IT reference genome; however, the indel error rate relative to the IT4 reference genome remained elevated. To correct small-scale errors and local misassemblies, we further polished this draft assembly with Illumina data using *Pilon V.1.22^15^*. The resultant hybrid Nanopore-Illumina assembly exhibited substantially less indel error, with 218 indels and 66 mismatches per 100kb (Table M1).

**Table M1:** Benchmarking of draft assemblies for BB12 against IT reference genome

|  | Canu only | Canu + Racon (x5) | Canu + Racon (x5) + Pilon |
| --- | --- | --- | --- |
| No. contigs | 17 | 17 | 17 |
| Total length (bp) | 21,817,649 | 22,819,515 | 23,097,724 |
| Largest contig (bp) | 3,142,754 | 3,312,166 | 3,350,589 |
| N50 length (bp) | 1,520,627 | 1,576,308 | 1,595,563 |
| N75 length (bp) | 1,298,640 | 1,358,305 | 1,375,631 |
| GC content (%) | 19.78 | 19.42 | 19.47 |
| Genome fraction (%) | 33.564 | 96.619 | 96.764 |
| Total aligned length | 7,539,785 | 22,142,137 | 22,497,519 |
| Mismatches (per 100kbp) | 89.10 | 150.97 | 65.84 |
| Indels (per 100 kbp) | 2216.89 | 1269.48 | 217.96 |

### Text S6: Gene Annotation and *var* Gene Domain Classification

Gene annotation of hybrid assemblies was performed with the *Companion* pipeline^16^ via the web server (https://companion.sanger.ac.uk) (July 2019 version), using *P. falciparum 3D7* as a reference strain and taking into account reference protein evidence. Highly conserved genes were mapped using the Rapid Annotation Transfer Tool (RATT), while a relatively lenient threshold inclusion score of 0.5 was used for de novo gene prediction by AUGUSTUS, to increase the sensitivity of gene annotation. Pseudogene detection was also performed as part of the Companion pipeline.

To characterise the ability of Nanopore long reads to access complex genomic regions, genes encoding either PfEMP1, DBL-domains, N-terminal or acidic-terminal segments (all associated with *var* genes) were identified. Protein sequences of all potential full and partial *var* genes were extracted and classified using the *VarDom 1.0 Server*^17^ (http://www.cbs.dtu.dk/services/VarDom/), with the default lower-bit score threshold 9.97 for homology blocks. Annotated *var* genes were considered to be complete if they included both acidic- and N-terminal segments. Accounting for the possible disruption of ORFs due to local assembly errors, neighbouring annotated *var* candidates within 1200bp windows were concatenated if their domain structures suggested they corresponded to the same gene to obtain ‘split’ *var* genes. *Var* candidates that included at least 3 annotated domains but did not contain either acidic- or N-terminal segments were also considered to be partial *var* genes.

### Text S7: Structural Variant Calling Pipelines

For short read data (Illumina), adapter trimming was first performed with *TrimGalore*. Trimmed Illumina reads were aligned against the 3D7 reference genome with *bwa mem V0.7.13*^2^, and resultant alignments files were sorted and indexed with *Samtools V1.7*^18^ utilities. Structural variant calling was then performed with *GRIDSS V2.8.3*^19^. Unpaired breakend variants were removed, and only simple variant types (i.e. insertions, deletions, duplications and inversions) were annotated using the R package *StructuralVariantAnnotation V1.0.0*.

Structural variants were retained if they passed default quality filters (i.e. the FILTER flag in the VCF files generated by GRIDSS was set to PASS).

For long read data (Nanopore), reads were first aligned against the 3D7 reference genome using *NGMLR V0.2.7*^20^ with the *ont* preset. The resultant alignments were sorted and indexed using *Samtools V1.7*^18^ utilities. Structural variant calling was subsequently performed with *Sniffles V1.0.11*^20^ using default parameters. Breakends were removed to retain only insertions, deletions, translocations, duplications and inversions. A threshold of 20 high-quality reads supporting the called variant (as determined by the RE tag in the output VCF file) was implemented to obtain high-quality structural variants. Overlapping duplications and deletions were assumed to be spurious and were accordingly removed.

To perform structural variant calling with our *de novo* assemblies, we first aligned our assemblies against 3D7 reference genome using *nucmer*(*MUMmer, V.3.1*^21^). All anchor matches were used, irrespective of their uniqueness, and a minimum cluster length of 500 was implemented; maximal exact matches were also required to be at least 100bp in length. The resultant delta files were processed using the Assemblytics web server^22^ to identify structural variants.

To quantify concordance rates across these methodologies we compared the set of high-quality SVs detected by each pipline using a custom *R* script. SVs identified by different pipelines were considered to be equivalent if: (i) they overlapped by at least 1bp; (ii) were annotated within the same SV class (Table M2); and (iii) if the length of the shorter SV was at least 75% of the length of the longer SV. To account for systematic indel errors in MinION basecalling, particularly in homopolymeric tracts, we retained only SVs with length at least 200 bp^20^. SVs exceeding 300 kbp seemed to be spurious across all three pipelines and were thus removed.

**Table M2:**
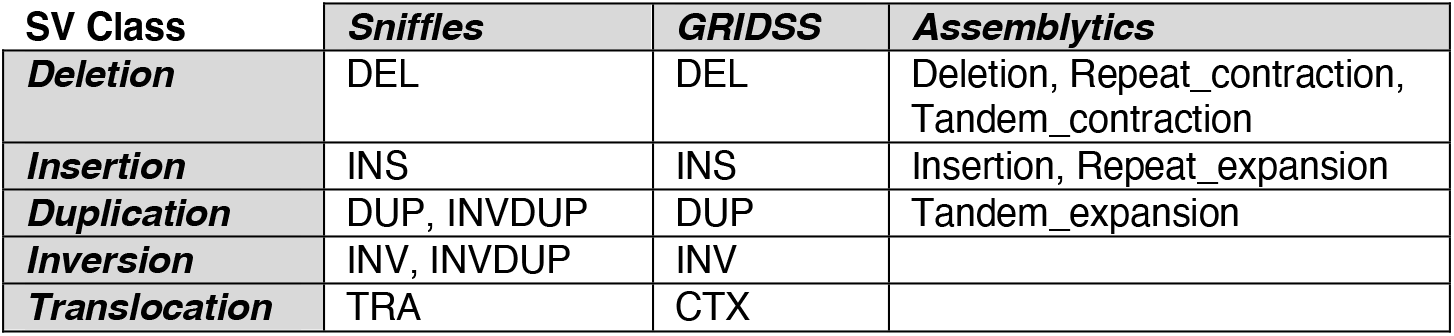
SV classes identified by each pipeline

## SUPPORTING FIGURES

**Figure S1.** Optimised bioinformatics pipelines

**Figure S2.** Parameters associated with Nanopore sequencing output, by flow cell

**Figure S3.** Relative frequencies of concordant and discordant genotype calls using Nanopore and Illumina data

**Figure S4.** Alignments of de novo BB12 and XHA assemblies against reference genomes

**Figure S5.** Domain structures for annotated *var* genes

**Figure S1A.**
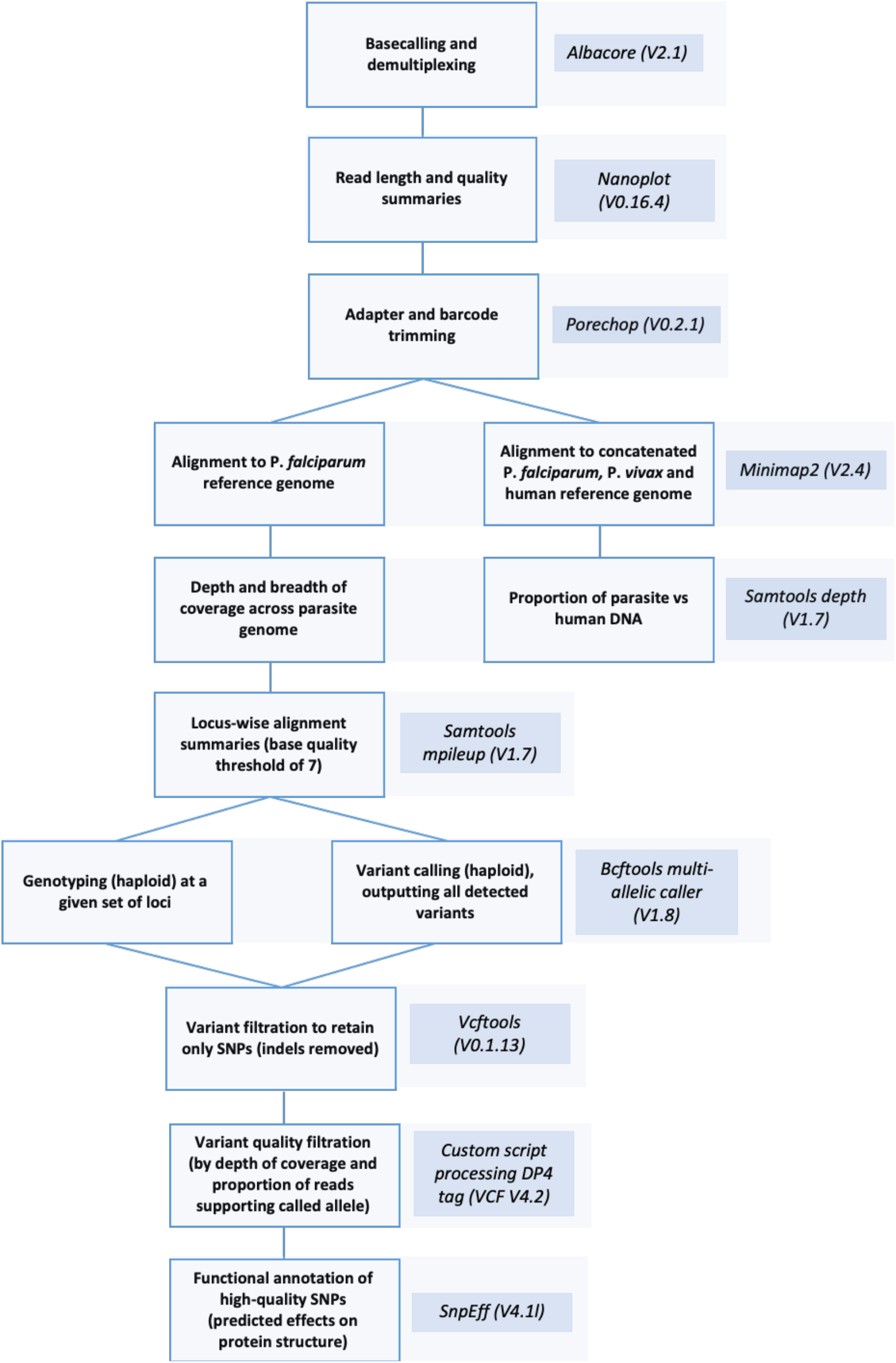
Variant calling and alignment pipeline for Nanopore data

**Figure S1B.**
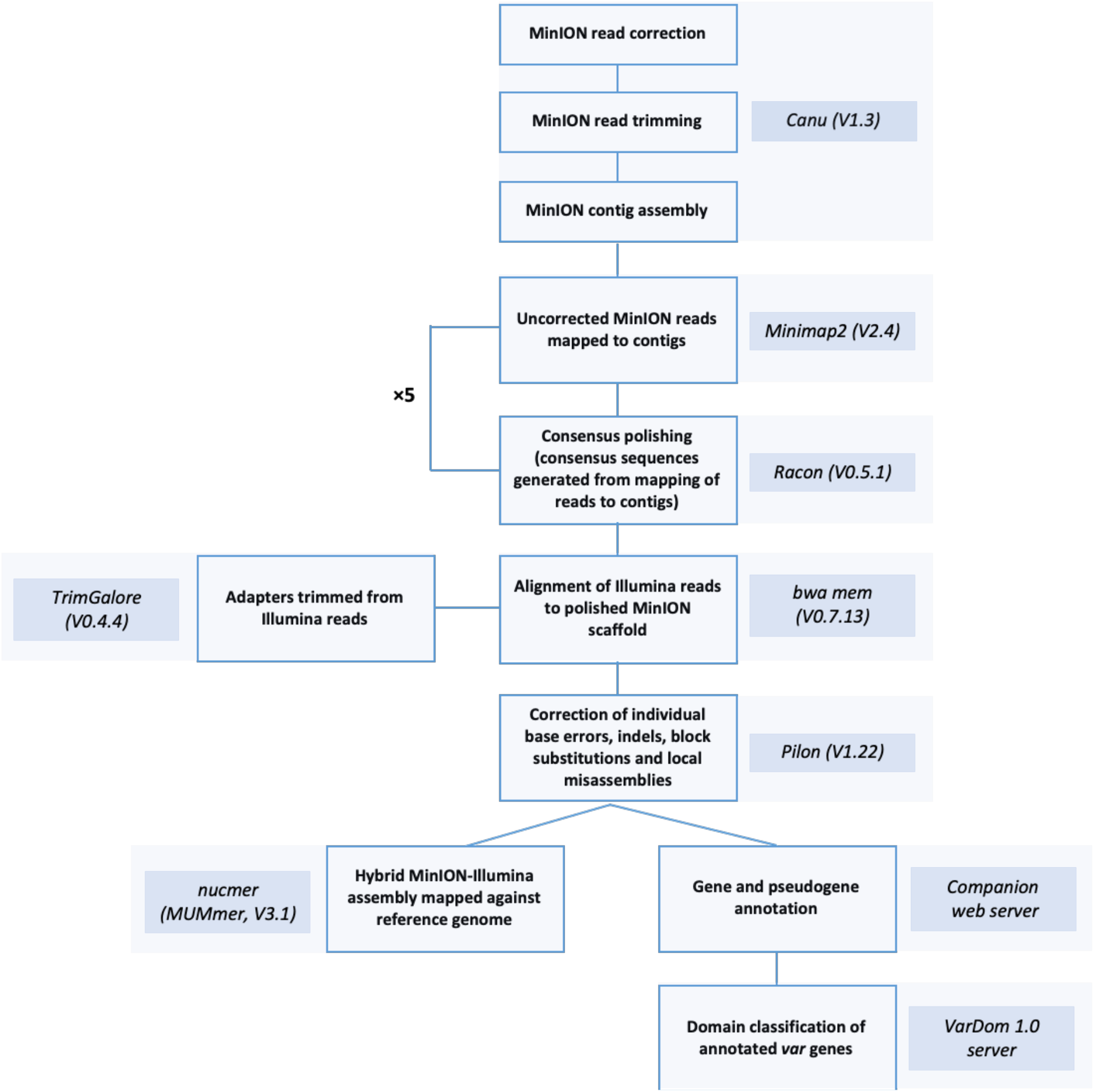
*De novo* assembly pipeline for hybrid assemblies of Nanopore and Illumina data

**Figure S2.**
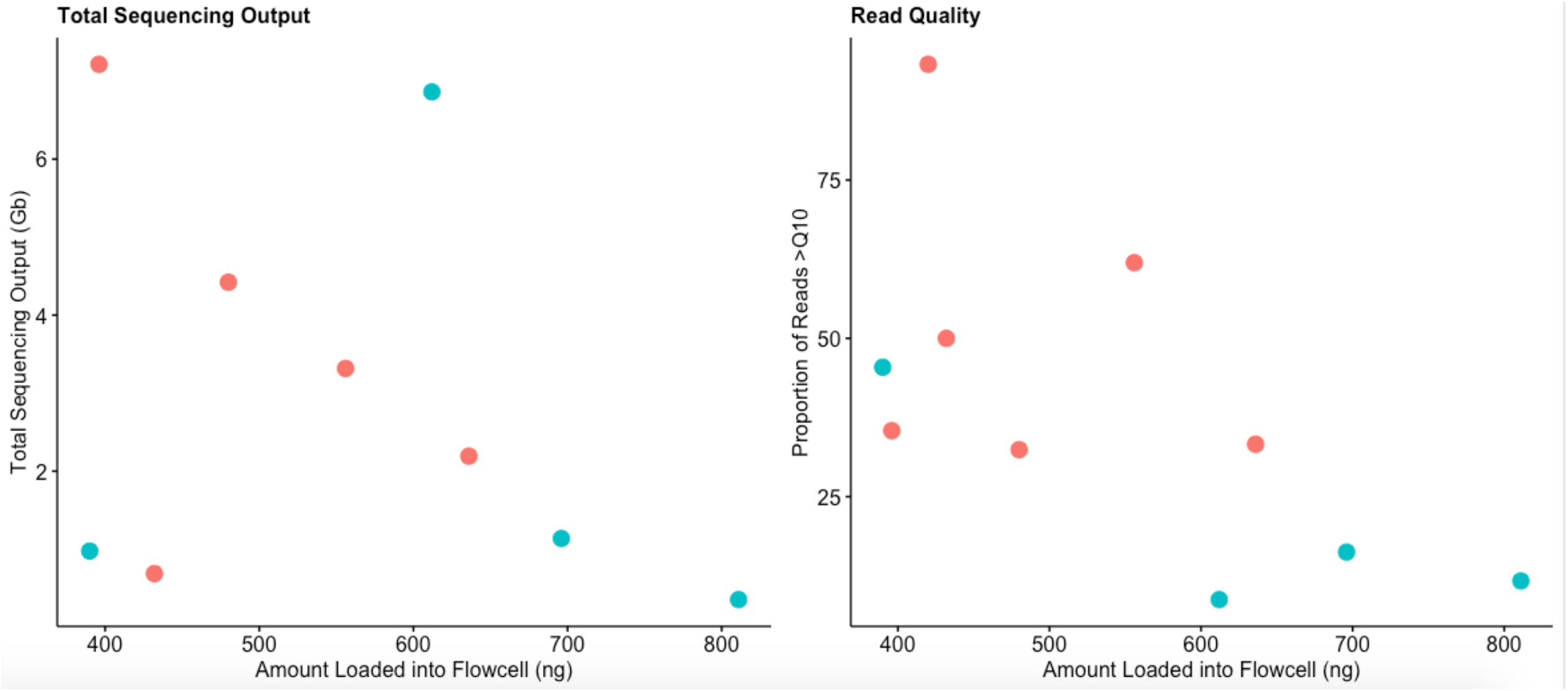
Parameters associated with Nanopore sequencing output, by flowcell. Nanopores tend to become saturated when large amounts of DNA are loaded into a flowcell, leading to a generally decreasing trend between the total sequencing output/quality and the amount of loaded DNA.

**Figure S3.**
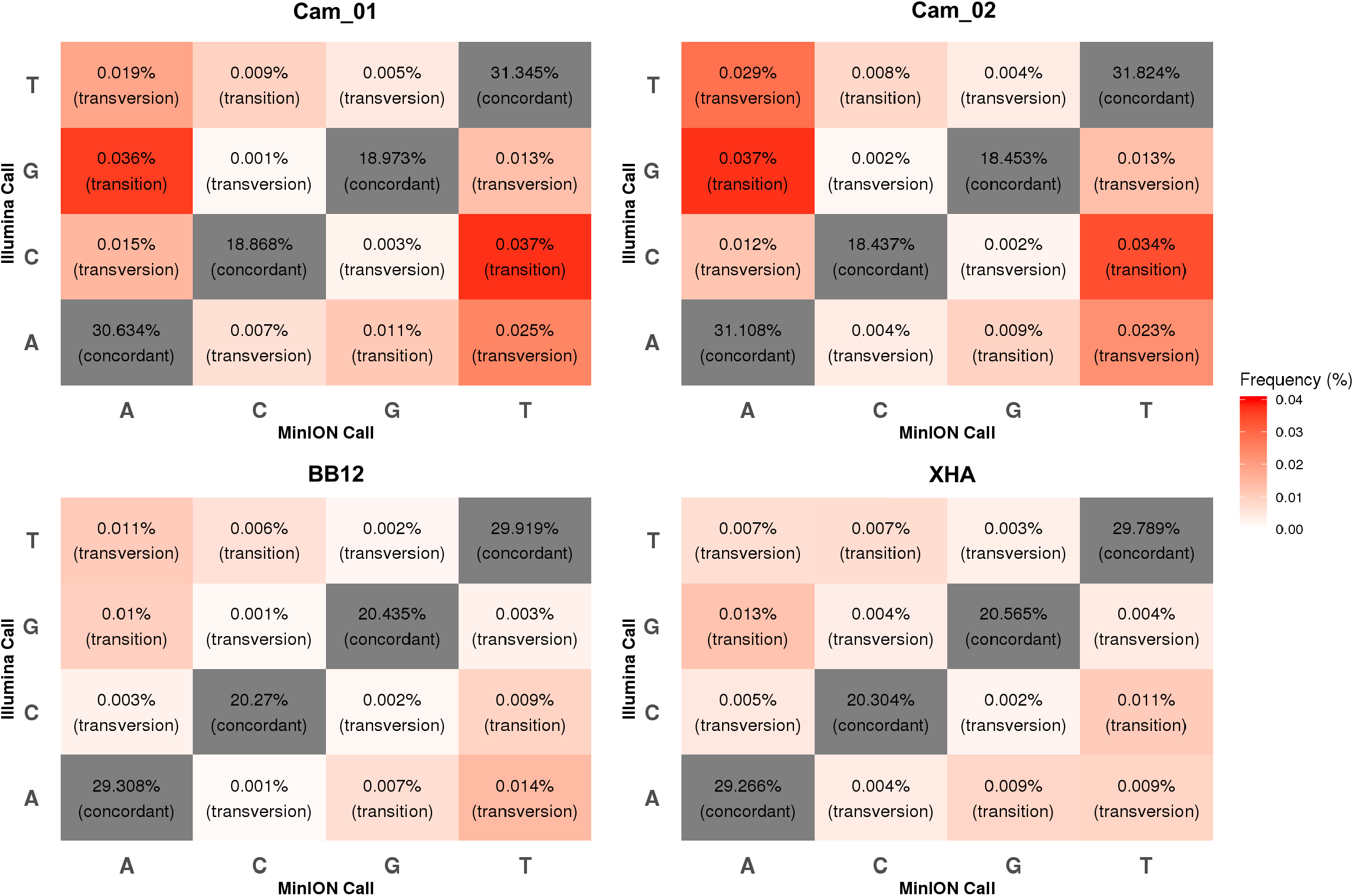
Relative frequencies of concordant and discordant genotype calls using Nanopore and Illumina data. Discordant calls were stratified by transition/transversion. Transition errors (between bases with similar ring structures) tend to be more common than transversion errors.

**Figure S4.**
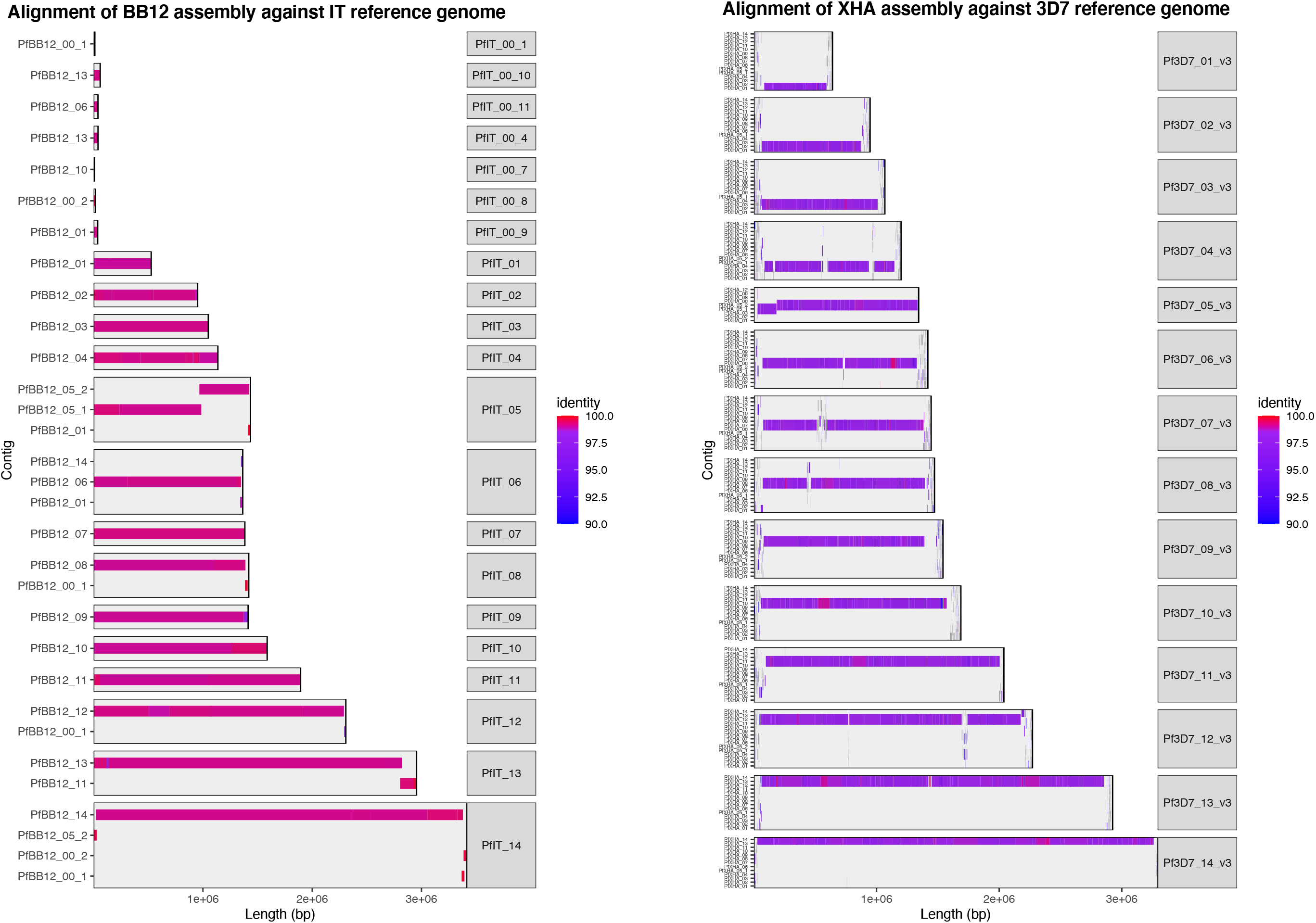
Alignments of *de novo* BB12 and XHA assemblies against reference genomes. IT4 (version 4) and 3D7 (version 3) respectively, with one-to-one mappings between de novo contigs and reference chromosomes computed using nucmer (MUMmer, V.3.1). Assembly continuity is high, with all but one nuclear chromosome spanned by a single contig. De novo contigs and reference genomes are generally concordant in core genomic regions, but alignments tend to become fragmented in subtelomeric hypervariable region

**Figure S5.**
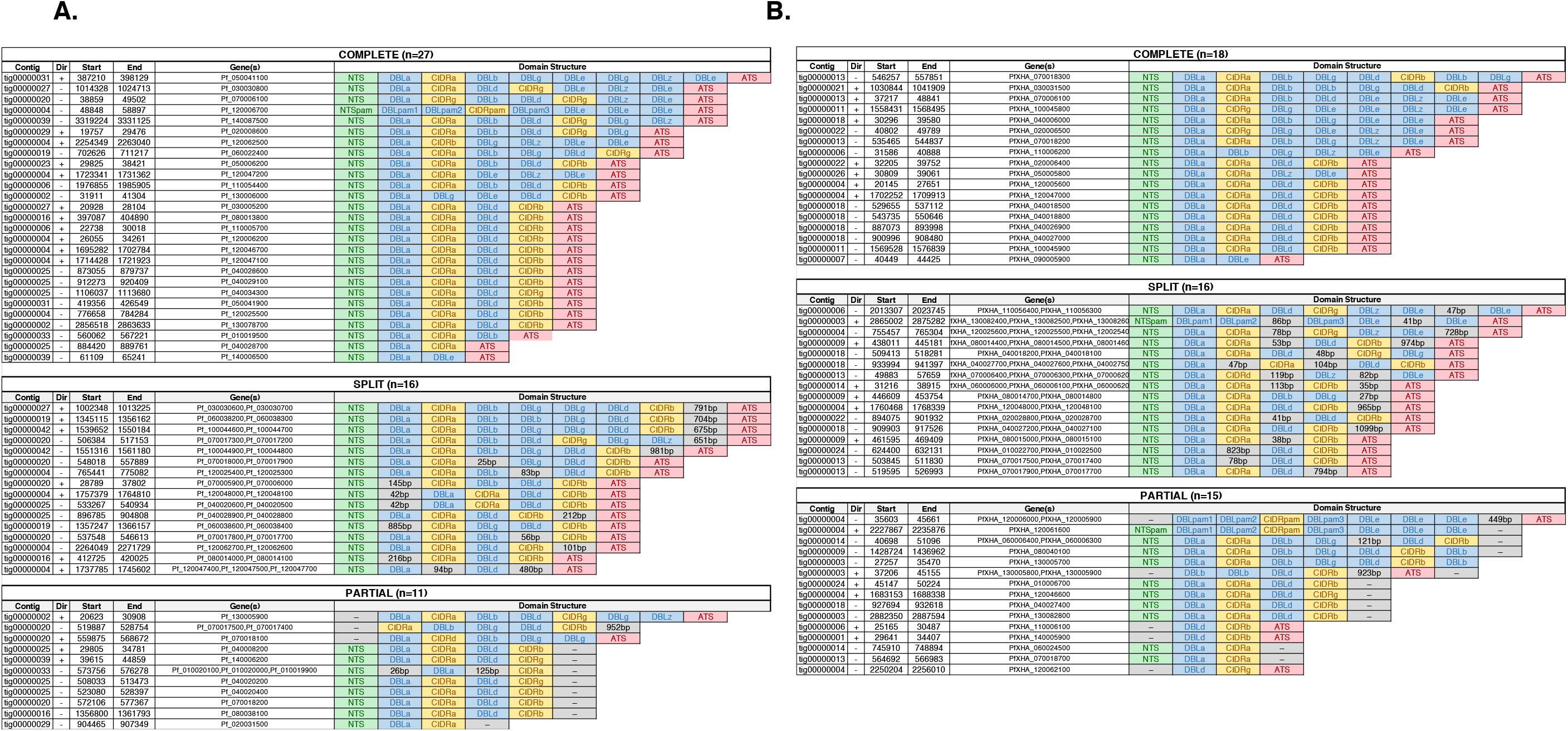
Domain structures for annotated *var* genes. *Var* gene structures were identified after *de novo* assembly of reads for *P. falciparum* laboratory isolates A) BB12 B) XHA

## SUPPORTING TABLES

**Table S1.** Primers and probes used in duplex qPCR assay

**Table S2.** Results of sequencing experiments

**Table S3**. SNV genotyping concordance rates for two laboratory and two field isolates in homopolymeric regions

**Table S1.**
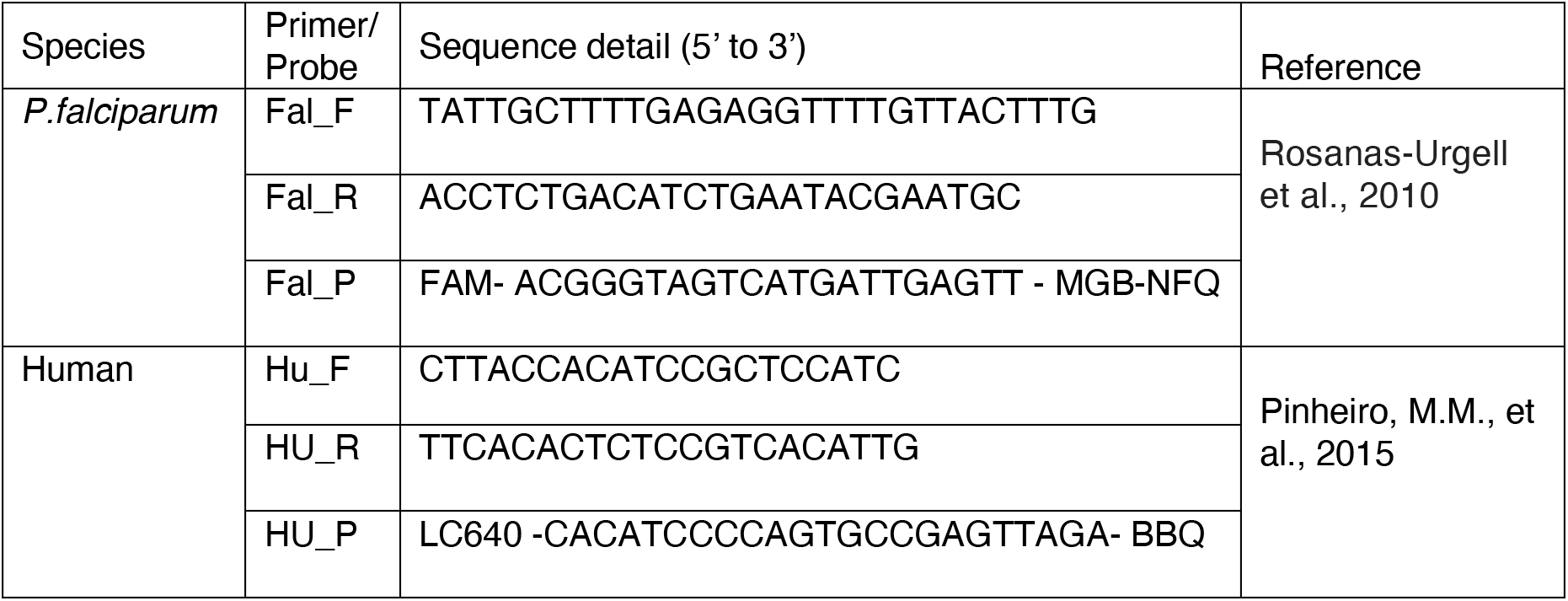
Primers and Probes used in Duplex qPCR Assay

**Table S2A.** Nanopore Sequencing Output for Pure Cultured Lines

| Sample ID | Sequencing Output |  |  |  |  |  | Depth and Breadth of Coverage (Pf Genome) |  |  |  |  |  |  |  |  |  |  |  | Read Qualities |  |  |
| --- | --- | --- | --- | --- | --- | --- | --- | --- | --- | --- | --- | --- | --- | --- | --- | --- | --- | --- | --- | --- | --- |
|  | Total bases (Gb) | Total reads | Mean read length (kb) | Median read length (kb) | Longest read length (kb) | Read length N50 (kb) | Mean depth of coverage | 1x breadth of coverage | 5x breadth of coverage | 10x breadth of coverage | 20x breadth of coverage | 30x breadth of coverage | 50x breadth of coverage | 75x breadth of coverage | 100x breadth of coverage | 150x breadth of coverage | 200x breadth of coverage | 250x breadth of coverage | Proportion >Q5 | Proportion >Q10 | Proportion >Q15 |
| XHA | 3.42 | 388,561 | 8.80 | 4.58 | 261.32 | 14.63 | 120.30 | 99.38% | 98.67% | 98.21% | 97.26% | 96.51% | 95.09% | 92.88% | 86.91% | 5.38% | 1.53% | 0.95% | 99% | 76% | 16% |
| BB12 | 2.07 | 298,286 | 6.95 | 3.44 | 157.32 | 15.26 | 76.06 | 99.55% | 98.85% | 98.08% | 96.46% | 94.63% | 89.90% | 54.85% | 5.50% | 1.54% | 1.14% | 0.70% | 92% | 69% | 0% |

**Table S2B.**
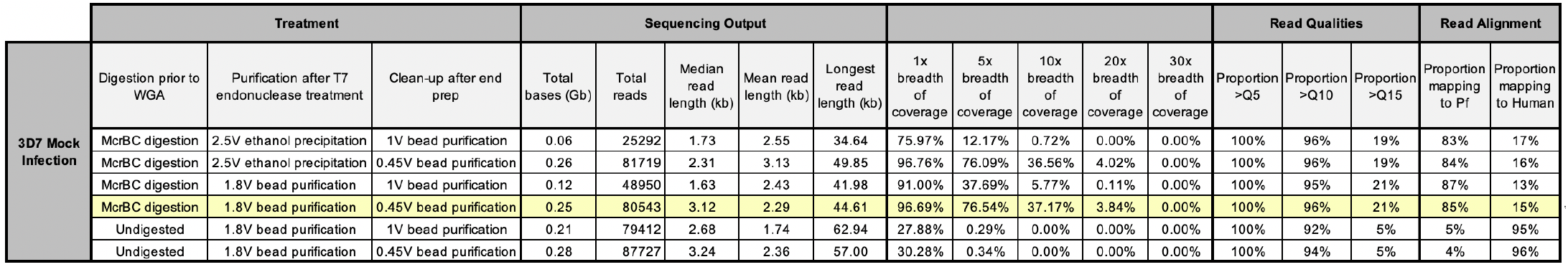
Nanopore Sequencing Output for Mock Infections

**Table S2C.**
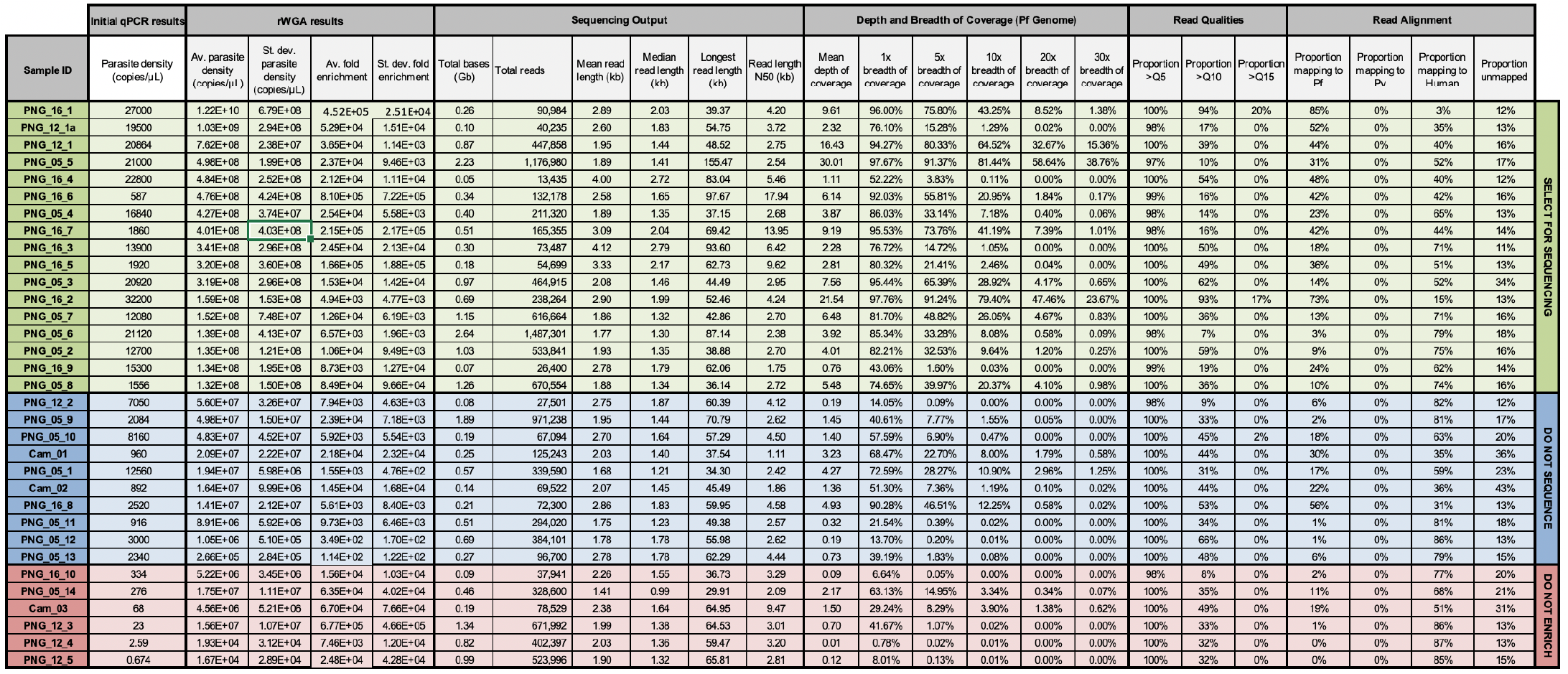
Nanopore Sequencing Output for Field Isolates

**Table S3.** SNV Genotyping Concordance Rates in Homopolymeric Regions

**Sample: XHA (106X coverage)**
|  | *Genotype* | *Unfiltered* | *Filtered* |
| --- | --- | --- | --- |
| Homopolymer <=6bp | Concordant | 683731 | 676846 |
|  | Discordant | 1649 | 507 |
| Homopolymer >6bp | Concordant | 7823 | 7641 |
|  | Discordant | 106 | 19 |
| *Discordance in homopolymers <=6bp:* |  | 0.24% | 0.08% |
| *Discordance in homopolymers >6bp* |  | 1.34% | 0.25% |

**Sample: BB12 (75X coverage)**
|  | *Genotype* | *Unfiltered* | *Filtered* |
| --- | --- | --- | --- |
| Homopolymer <=6bp | Concordant | 656892 | 649945 |
|  | Discordant | 1179 | 399 |
| Homopolymer >6bp | Concordant | 7560 | 7475 |
|  | Discordant | 99 | 54 |
| *Discordance in homopolymers <=6bp:* |  | 0.18% | 0.06% |
| *Discordance in homopolymers >6bp* |  | 1.29% | 0.72% |

**Sample: Cam\_01 (3.2X coverage)**
|  | *Genotype* | *Unfiltered* | *Filtered* |
| --- | --- | --- | --- |
| Homopolymer <=6bp | Concordant | 451550 | 303559 |
|  | Discordant | 3528 | 529 |
| Homopolymer >6bp | Concordant | 5076 | 3170 |
|  | Discordant | 161 | 27 |
| *Discordance in homopolymers <=6bp:* |  | 0.78% | 0.17% |
| *Discordance in homopolymers >6bp* |  | 3.07% | 0.84% |

**Sample: Cam\_02 (1.4X coverage)**
|  | *Genotype* | *Unfiltered* | *Filtered* |
| --- | --- | --- | --- |
| Homopolymer <=6bp | Concordant | 329574 | 169799 |
|  | Discordant | 3083 | 282 |
| Homopolymer >6bp | Concordant | 3628 | 1753 |
|  | Discordant | 146 | 23 |
| *Discordance in homopolymers <=6bp:* |  | 0.93% | 0.17% |
| *Discordance in homopolymers >6bp* |  | 3.87% | 1.30% |

## Notes

### Competing Interest Statement

The authors have declared no competing interest.

